# dAux orchestrates the phosphorylation-dependent assembly of the lysosomal V-ATPase in glia and contributes to α-synuclein degradation

**DOI:** 10.1101/2023.12.07.570521

**Authors:** Shiping Zhang, Linfang Wang, Shuanglong Yi, Yu-ting Tsai, Honglei Wang, Shuhua Li, Ruiqi Wang, Yang Liu, Wei Yan, Chang Liu, Kai-Wen He, Margaret S. Ho

## Abstract

Glia serve as double-edged swords to modulate neuropathology in Parkinson’s disease (PD), but how they react opposingly to be beneficial or detrimental under pathological conditions, like promoting or eliminating α-synuclein (α-syn) inclusions, remains elusive. Here we present evidence that dAuxilin (dAux), the *Drosophila* homolog of the PD risk factor Cyclin G-associated kinase (GAK), regulates the lysosomal degradation of α-syn in glia. Lack of glial Gak/dAux increases the lysosome number and size, regulates lysosomal acidification and hydrolase activity, and ultimately blocks the degradation of substrates including α-syn. Whereas α-syn accumulated prominently in lysosomes devoid of glial dAux, levels of injected α-syn preformed fibrils also further enhanced in the absence of microglial Gak. Mechanistically, dAux mediates phosphorylation at the serine 543 of Vha44, the V1C subunit of the vacuolar H^+^-ATPase (V-ATPase), regulates its assembly to control proper acidification of the lysosomal milieu. Expression of Vha44, but not the Vha44 variant lacking S543 phosphorylation, restores lysosome acidity, locomotor deficits, and DA neurodegeneration upon glial dAux depletion, linking this pathway to PD. Our findings identify a phosphorylation-dependent switch controlling the V-ATPase assembly for lysosomal α-syn degradation in glia. Targeting the clearance of glial α-syn inclusions via this lysosomal pathway could potentially be a therapeutical approach to ameliorate the disease progression in PD.

## Introduction

Protein inclusions in the brains are the histological hallmarks for neurodegenerative diseases, and a strategic way to target them for elimination is urgently needed to resolve the pathology. Whereas Lewy bodies (LBs), the fibrillar aggregates of α-synuclein (α-syn), accumulate in neurons in Parkinson’s disease (PD), evident α-syn inclusions are also detected in microglia, astrocytes, and oligodendrocytes due to prion-like propagation from neurons to glia^1–4^. In addition, glia (astrocytes and microglia) are double-edged swords in PD and respond to signals from neuronally-secreted α-syn by releasing pro- or anti-inflammatory factors to enhance or ameliorate PD progression, respectively^5–10^. While it is clear that α-syn inclusions in neurons are eliminated via the autophagy-lysosome pathway, the mechanism of how α-syn inclusions are eliminated in glia remains largely elusive.

Lysosomes, the acidic milieu containing hydrolases and other enzymes, are the final destinations for endocytic and autophagic pathways to digest extracellular and intracellular materials^11–13^. To maintain an optimal pH, the vacuolar H^+^-ATPase (V-ATPase) localizes on the lysosomal membrane and pumps protons into the lumen^14–16^. V-ATPase is a multisubunit enzyme composed of a cytosolic V1 domain with eight subunits (A-H) mediating ATP hydrolysis and a membrane-bound V0 domain with six subunits (a, c, c”, d, e, and c’ in yeast or Ac45 in higher eukaryotes) mediating proton transport^16–18^. Using the energy generated by ATP hydrolysis, V-ATPase drives the ion flux to regulate a variety of cellular processes, such as endosome maturation and trafficking, receptor recycling, and nutrient signaling^19, 20^. Notably, the function of V-ATPase in controlling lysosomal acidification could well be a key factor tuning the level of protein inclusions in the diseased brains; how V-ATPase contributes to the pathological aggregate formation and elimination, awaits to be further explored.

Here we present evidence for a lysosomal clearance pathway of α-syn degradation in glia. dAuxilin (dAux), the *Drosophila* homolog of the human Cyclin G-associated kinase (GAK, mouse Gak), mediates the serine 543 (S543) phosphorylation of Vha44, the catalytic C subunit of the lysosomal V-ATPase V1 complex. Lack of dAux decreases Vha44 S543 phosphorylation, reduces V-ATPase assembly, and disrupts lysosomal acidification and hydrolase activity in glia. These defects ultimately cause α-syn accumulation, leading to a broad spectrum of PD-like symptoms including DA neurodegeneration and locomotor deficits in flies and mice. Our findings suggest that targeting the lysosomal α-syn degradation in glia might be a strategic way to eliminate protein inclusions in PD.

## Results

### Lack of dAux increases the lysosome number in adult fly glia

Our recent study indicated that Gak/dAux regulates autophagy initiation in glia^21, 22^. Promoting autophagy initiation, however, is in apparent contradiction to the observed block of substrate degradation in the absence of Gak/dAux, leading us to speculate whether Gak/dAux confers additional regulations. We then further investigated the cellular effect of lacking dAux in adult fly glia. As previously reported, dAux expression in glia was manipulated by two independent transgenic fly lines: *UAS-daux*-RNAi (*daux*-RNAi, V#16182) or *UAS-daux*-RNAi^#2^ (*daux*-RNAi^#2^, BL#39017) under the control of the widely used pan-glial driver *repo*-GAL4 (*repo*>*daux*-RNAi or *daux*-RNAi^#2^). Upon RNAi expression, dAux levels were reduced by about 50%, validating the RNAi efficiency^21^. For comparison of the RNAi effect, *UAS-LacZ* (*LacZ*) was used as a control throughout the study, while UAS-*luciferase*-RNAi (*luc*-RNAi) was used in some experiments, with similar conclusions.

Given the role of Gak/dAux in autophagy and its implication in PD, we asked if dAux is involved in the lysosomal degradation pathway in adult fly glia, as lysosomes are the terminal milieu converged from multiple pathways for degradation of substrates like α-syn. In these experiments, a selected anterior/dorsal glia-rich region in the adult fly brain was analyzed with glial lysosomes labeled by *repo*>*Lamp1-GFP* (Figures 1A and 1B). Interestingly, the number of glial Lamp1-positive lysosomes increased in the 10-day-old adult fly brains upon *daux*-RNAi expression (Figures 1C and 1D). Results were consistent when comparing with the *luc*-RNAi or using the *daux*-RNAi^#2^ (Figures S1A-S1D). Some of these lysosomes were enlarged and clustered, implying non-degraded substrate accumulation (Figures 1C, 1D, and S1A-S1D)^23–25^. Ultrastructural analysis by Transmission Electron Microscopy (TEM) also revealed an increase in the lysosome number and size in the absence of glial dAux (Figures 1E-1G, yellow arrowheads). Notably, some of the large lysosomes exhibited electron-lucent dots, suggesting disrupted lysosomal pH^26, 27^ (Figures 1F and 1F’, red arrowheads). Of note, the overall brain size and perimeter of flies lacking glial dAux were larger and longer, respectively (Figures S1E and S1F). Yet, the numbers of total glial cells, Rab5-positive early endosomes, and Rab7-positive late endosomes remain largely unaffected in the absence of glial dAux^21^. Taken together, these results suggest that dAux regulates the lysosome number in glia.

**Figure 1.**
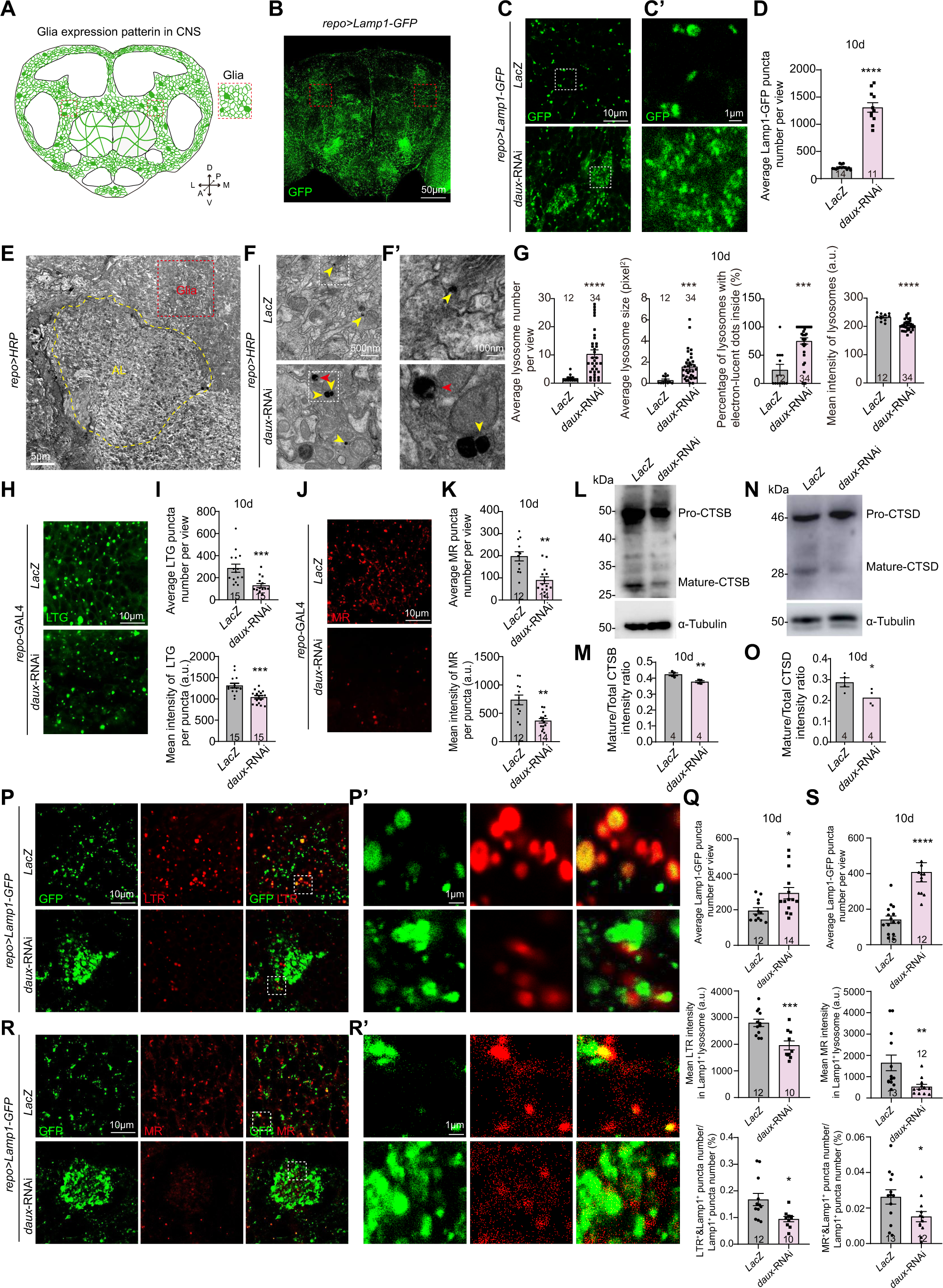

### Lack of Gak increases the lysosome number in immortalized microglia and mouse primary microglia

To further demonstrate if the dAux-mediated increase in the lysosome number is conserved across species, we investigated the cellular consequence of lacking the mammalian *daux* homolog, *Gak*. Given that microglia are major mediators of PD and Gak expression is relatively higher in microglia^28, 29^, we inhibited Gak expression in the mouse immortalized microglial cell line (IMG)^30^ and primary mouse microglia. The IMG identity, *Gak* siRNA (*Gak* siRNA^#1^ and *Gak* siRNA^#2^) knock-down efficiency in IMG, and the *Gak* expression level in the mouse conditional knockout (cKO) microglia (*CX3CR1-Cre^+/−^; Gak^flox/flox^*) have been validated by antibody staining, western blot (WB), and qRT-PCR analyses, demonstrating the reduced Gak level in both systems^21^. Consistent with our results in adult fly glia, the lysosome intensities in both siRNA-treated IMG and *Gak* cKO microglia were elevated (Figures S2A-S2D). The Lamp1-positive puncta size and the colocalization of Lamp1 and ionized calcium-binding adapter molecule 1 (Iba1) in *Gak* cKO microglia were also increased (Figures S2C and S2D). Taken together, these results suggest that fly dAux function on lysosome biogenesis is conserved in mammalian microglia.

### Lack of dAux disrupts lysosomal acidification in adult fly glia

We next asked if lysosome function is affected by the absence of glial dAux. To address this question, Lysotracker Green (LTG), a fluorescent dye that specifically labels acidic organelles including lysosomes, was used to probe lysosome pH. Interestingly, the number and intensities of glial LTG-positive puncta were significantly reduced in the 10-day-old adult fly brains lacking dAux, indicating that lysosomes are not properly acidified (Figures 1H, 1I, and S3A-S3D). The activity of the lysosomal hydrolase Cathepsin B (CTSB) stained by Magic Red (MR) was also significantly decreased (Figures 1J, 1K, and S3E-S3H). In addition, the processing of CTSB and another lysosome hydrolase Cathepsin D (CTSD) was impaired, suggesting a general deficit in the lysosome hydrolase activity (Figures 1L-1O). In-depth analysis revealed that the number of glial Lamp1-positive lysosomes increased, while the mean intensities of Lysotracker Red (LTR) or MR per glial Lamp1-positive lysosome and the percentage of the number for LTR-Lamp1 co-positive puncta over total glial Lamp1-positive lysosomes puncta (LTR^+^&Lamp1^+^ puncta number/Lamp1^+^ puncta number, %) decreased significantly upon dAux depletion (Figures 1P-1S and S3I-S3N). It is noteworthy to mention that the expression of either fly dAux or human GAK (Flag-hGAK) fully rescued the *daux*-RNAi-induced lysosomal acidification defects^21^ (Figures S3O and S3P), reinforcing the conserved function for GAK/dAux across species. Furthermore, the intensities of LTG- or LTR-positive puncta remain largely unaffected in the whole brain or per glial Lamp-1 positive lysosome, respectively when expressing *daux*-RNAi using a pan-neuronal driver *GMR57C10*-GAL4, suggesting that dAux exhibits minimum effect on lysosomal acidification in neurons (Figures S4A-S4D). In summary, lack of dAux disrupts lysosomal acidification and hydrolase activity in glia.

### Lack of glial dAux causes accumulation of Ref(2)P, Ubi, and **α**-syn

To investigate the consequence of impaired lysosomal acidification, we analyzed the levels of known substrates to reflect the efficiency of lysosomal degradation. Interestingly, autophagic substrates Ref(2)P (the *Drosophila* ortholog of P62) and Ubiquitin (Ubi) accumulated in the absence of glial dAux (Figures 2A, 2B, and S5A-S5D). In addition, a binary system (GAL4/lexA) that allows simultaneous expression in neurons and glia was established. Upon co-expression of human α-syn in neurons and *daux*-RNAi in glia, total α-syn levels in the adult fly brains increased, suggesting that α-syn accumulates in the absence of glial dAux (Figures 2C and 2D). Further in-depth analysis showed that the accumulated α-syn in adult fly brains were enriched in glial lysosomes as both the α-syn intensities inside the Lamp1-positive lysosomes and α-syn-Lamp1 colocalization increased upon *daux*-RNAi expression in glia (Figures 2E and 2F). Taken together, these results suggest that dAux mediates the lysosomal degradation of α-syn and other substrates in glia.

**Figure 2.**
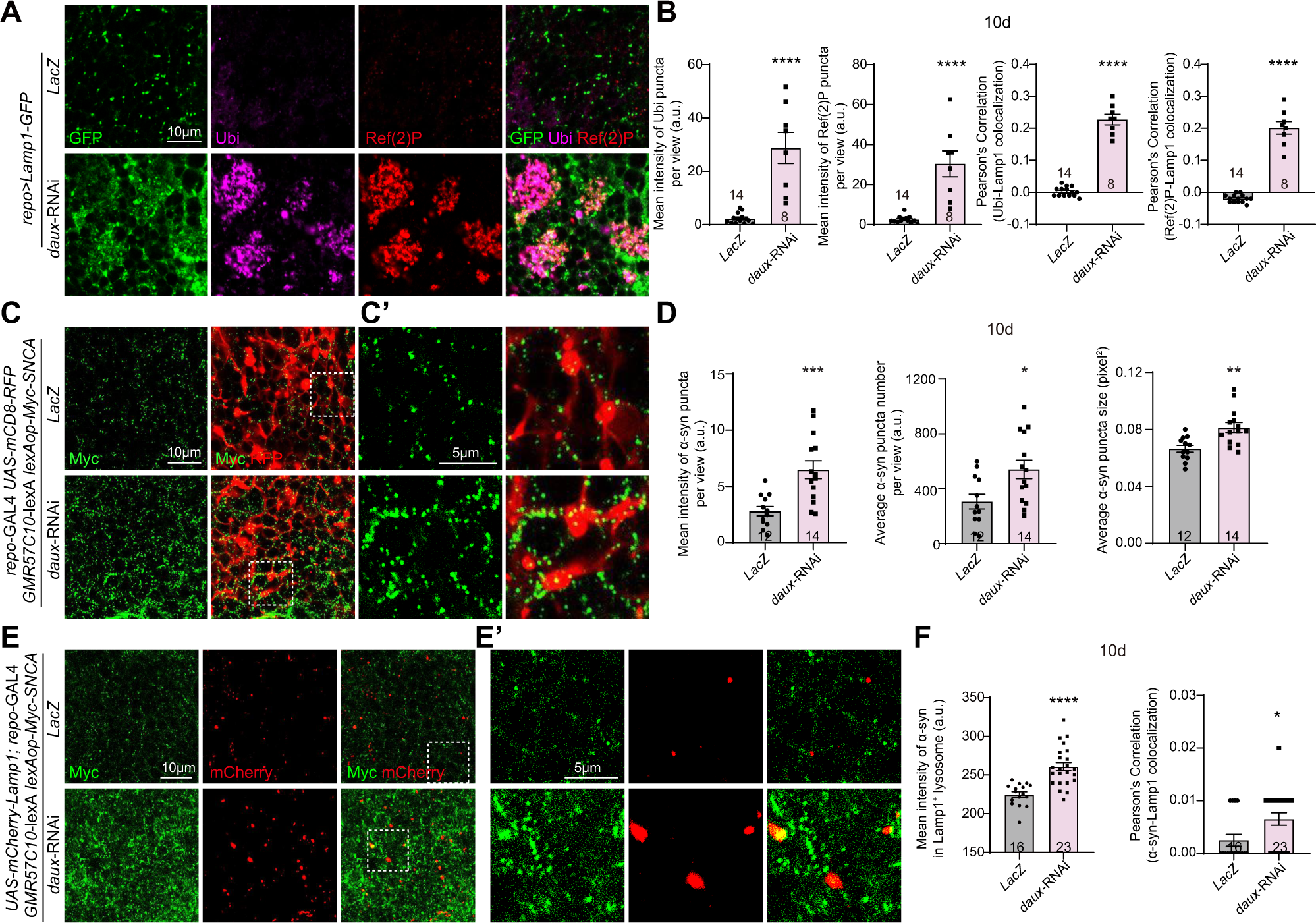

### dAux regulates Vha44 S543 phosphorylation

Previous studies have revealed an impact on the lysosome homeostasis by GAK kinase inhibitors^23, 24^. However, whether dAux regulates the lysosome number and acidification via its kinase activity remains unexplored. In our rescue experiments, expression of the dAux variant lacking the kinase domain (Flag-dAux^ΔKinase^)^21^ failed to restore the *daux*-RNAi-induced lysosomal acidification defects, suggesting that dAux kinase activity might contribute (Figures S3O and S3P). To this end, phosphoproteomic analysis detecting changes in phosphorylated peptides was conducted using 10-day-old adult fly heads of the control and *repo>daux*-RNAi flies^21^ (Figures S6A-S6E). Interestingly, a list of proteins encoding different subunits of the lysosomal V-ATPase, the main driving force in maintaining the lysosome pH, were identified (Figures 3A-3D). Among all, the phosphorylation level at the S543 of the V-ATPase V1C subunit, Vha44 isoform F, was significantly decreased upon glial dAux depletion in two out of three replicates (*daux-*RNAi/LacZ ratio 0.634, Figure 3D). Motif analysis also predicted that S543 belongs to a sequence motif prone to be phosphorylated (xxxxPx_S_Pxxxxx, Figure 3E). The phosphorylation level at the Y112 of another subunit Vha26 appears to be consistently increased (*daux-*RNAi/LacZ ratio 1.484, Figures 3C and 3D). Given that we are looking for substrates with reduced phosphorylation upon glial dAux depletion, we focused on the analysis of Vha44 S543 phosphorylation.

**Figure 3.**
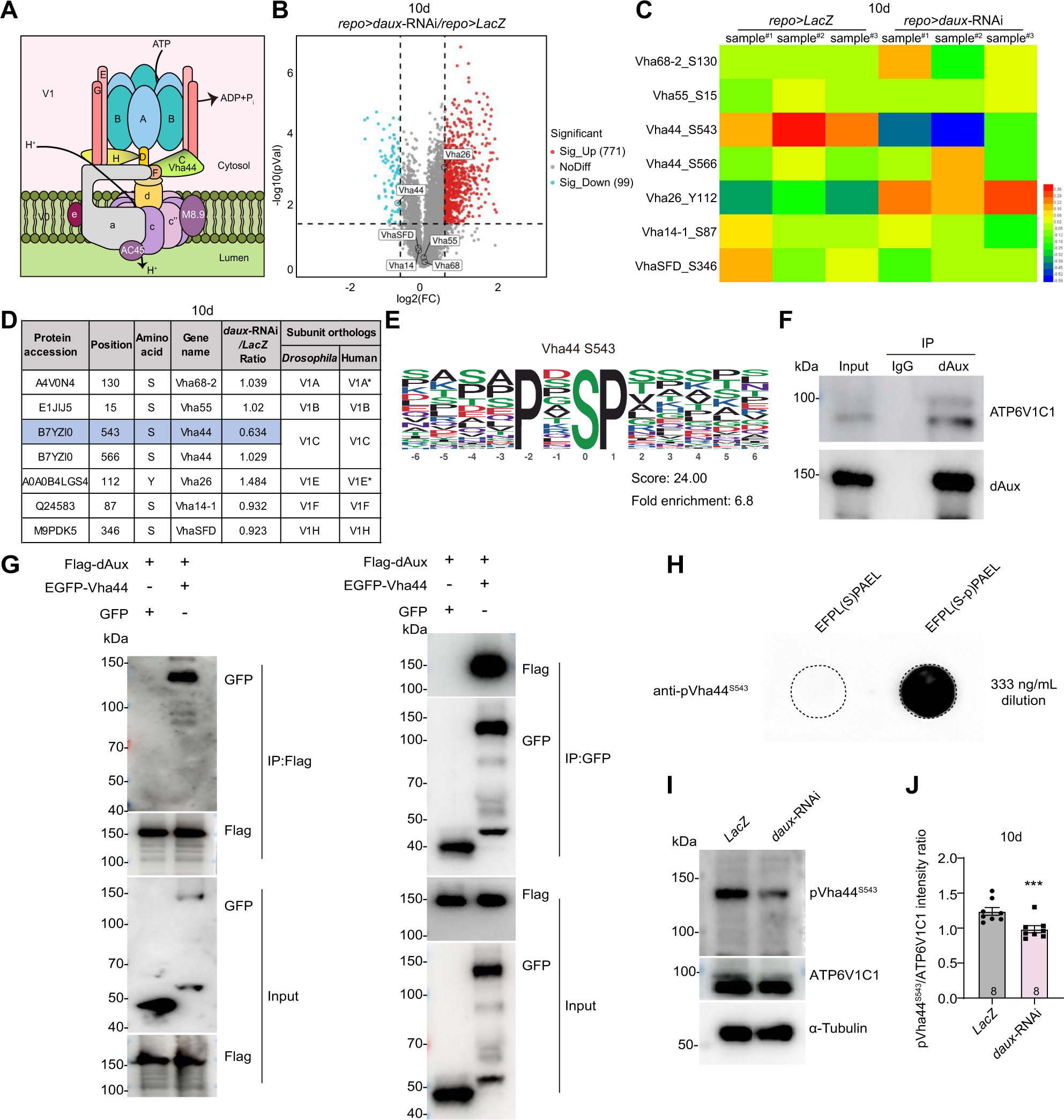

To support our hypothesis that Vha44 is a dAux kinase substrate, co-immunoprecipitation (Co-IP) analysis was first conducted. Our results revealed that dAux interacted with Vha44 both endogenously and upon overexpression in S2 cells (Figures 3F and 3G). In addition, antibodies that specifically recognize the synthesized phosphopeptides EFPL(S-p)PAEL, sequence correlating with the identified motif, were generated (anti-pVha44^S543^). Dot blot analysis validated the specificity of anti-pVha44^S543^ antibodies to the phosphopeptide immunogen (Figure 3H). Consistent with the results from the phosphoproteomic analysis, endogenous Vha44 S543 phosphorylation probed by the anti-pVha44^S543^ antibodies was significantly reduced in the 10-day-old adult fly brains expressing the *daux*-RNAi in glia (Figures 3I and 3J). These results suggest that dAux mediates Vha44 S543 phosphorylation.

### Vha44 regulates lysosomal acidification in glia

Next, we investigated if Vha44, like dAux, regulates lysosome acidification in glia. Consistent with the results from *daux*-RNAi-expressing flies, glial expression of *vha44*-RNAi, which efficiency was validated by qRT-PCR and WB analyses (Figures S7A-S7C), caused a reduction in the number of glial LTG- and MR-positive puncta in the 10-day-old adult fly brains (Figures 4A, 4B, S7D, and S7E). The intensities of the LTR-positive puncta per glial Lamp1-positive lysosome decreased significantly upon Vha44 depletion (Figures 4C and 4D). In contrast to dAux, the number of glial Lamp1-positive lysosomes decreased in the absence of Vha44, suggesting that Vha44 might regulate the lysosome number via different mechanisms (Figure 4D). Taken together, these results demonstrate that Vha44 regulates lysosomal acidification in glia.

**Figure 4.**
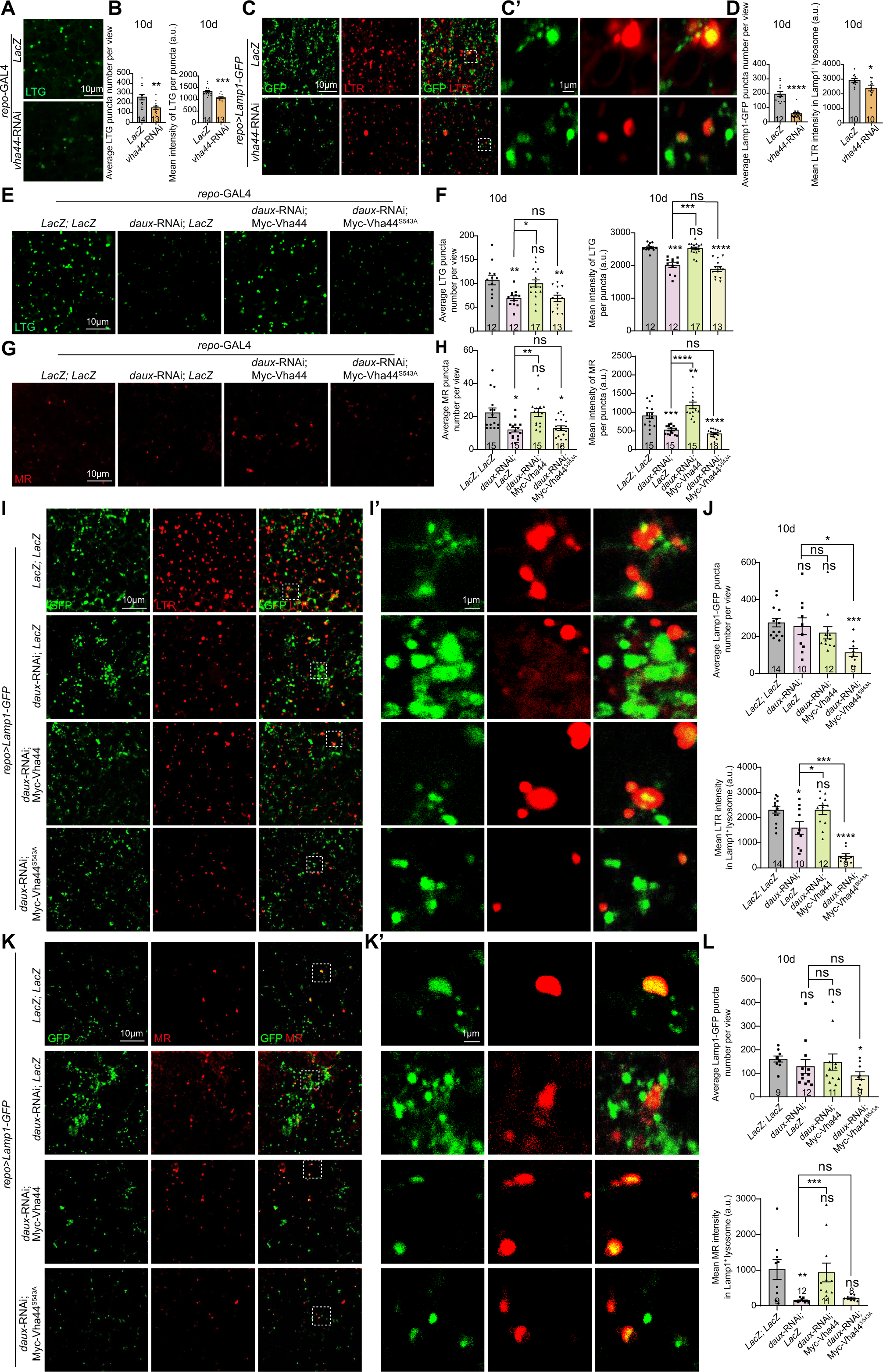

### dAux regulates lysosomal acidification and hydrolase activity via Vha44 S543 phosphorylation in glia

Given that both dAux and Vha44 regulate lysosomal acidification in glia, we next conducted epistatic analysis that helps to resolve the genetic relationship between dAux and Vha44 in the context of glial lysosomal acidification. Transgenic flies expressing the wild-type Vha44 (Myc-Vha44) or the non-phosphorylatable Vha44 mutant with alanine substitution of S543 (Myc-Vha44^S543A^) were generated. These Vha44 variants were expressed properly as validated by WB (Figure S7F). Notably, Myc-Vha44 expression significantly restored the *daux*-RNAi-mediated decrease in the intensities of LTG- or MR-positive puncta in the whole brain (Figures 4E-4H) and LTR- or MR-intensities per glial Lamp1-positive lysosome (Figures 4I-4L), respectively. Importantly, Myc-Vha44^S543A^ expression in glia failed to restore such decrease. Of note, the number of Lamp1-positive puncta in adult fly brains expressing *daux*-RNAi; *LacZ* exhibited a decreasing but not significant trend, potentially due to the titration effect of two *UAS* transgenes (Figures 4J and 4L). Taken together, these results suggest that Vha44 acts downstream of dAux and S543 phosphorylation is required for dAux-mediated lysosomal acidification in glia.

On the other hand, glial expression of Myc-Vha44^S543A^, but not Myc-Vha44, caused a significant reduction in the number and intensities of the LTG-positive puncta in the 10-day-old adult fly brains (Figures S7G and S7H). Furthermore, glial expression of either Myc-Vha44 or Myc-Vha44^S543A^ reduced the intensities of the MR-positive puncta in the brains, with the latter exhibiting a more potent suppression (Figures S7I and S7J). Taken together, these results suggest that supplying more of the non-phosphorylatable Vha44^S543A^ might disrupt the overall balance of the Vha44 protein pool, hence the dysregulation of dAux-mediated lysosomal acidification in glia.

### dAux-mediated Vha44 S543 phosphorylation regulates V-ATPase assembly in glia

V-ATPase is the key factor controlling lysosome pH by pumping protons into the lumen. As we saw consistent phenotypes on glial lysosomal acidification, we next asked if the V-ATPase function, mainly relying on its V1-V0 subcomplex assembly, is regulated by the dAux-mediated Vha44 S543 phosphorylation. Transgenic flies that express EGFP-Vha44 labeling the V1C subunit and mCherry-Vha100-2 labeling the V0a subunit were generated to analyze the fluorescent signal colocalization of these two subunits, reflecting how well the V-ATPase assembles (Figure 5A). As EGFP-Vha44 expression in glia revealed signals mainly distributed on the glial processes, mCherry-Vha100-2 expression in glia exhibited as puncta, localizing on the lysosomal membrane (Figure 5). Interestingly, the EGFP-mCherry co-positive signals, reflecting how well the two subunits colocalize, decreased significantly upon glial dAux depletion (Figures 5B and 5C). Using a different reporter mCherry-Lamp1, the EGFP-mCherry co-positive signals representing the Vha44 colocalization with Lamp1 also decreased significantly (Figures 5D and 5E). In addition, live-cell imaging analysis showed that the percentage of EGFP-positive puncta (Vha44) being trafficked to the vicinity of mCherry-positive puncta (Vha100-2) within a radius of 0.42 μm decreased (Figures 5F and 5G) and the duration time of the Vha44-Vha100-2 contact was shorter (Figures 5F and 5H) upon glial dAux depletion. The trafficking speed of EGFP- and mCherry-positive puncta increased and remained unaffected, respectively in the absence of glial dAux, suggesting that Vha44 is more mobile (Figures 5F and 5I). Altogether, these results indicate that glial dAux regulates the V-ATPase assembly.

**Figure 5.**
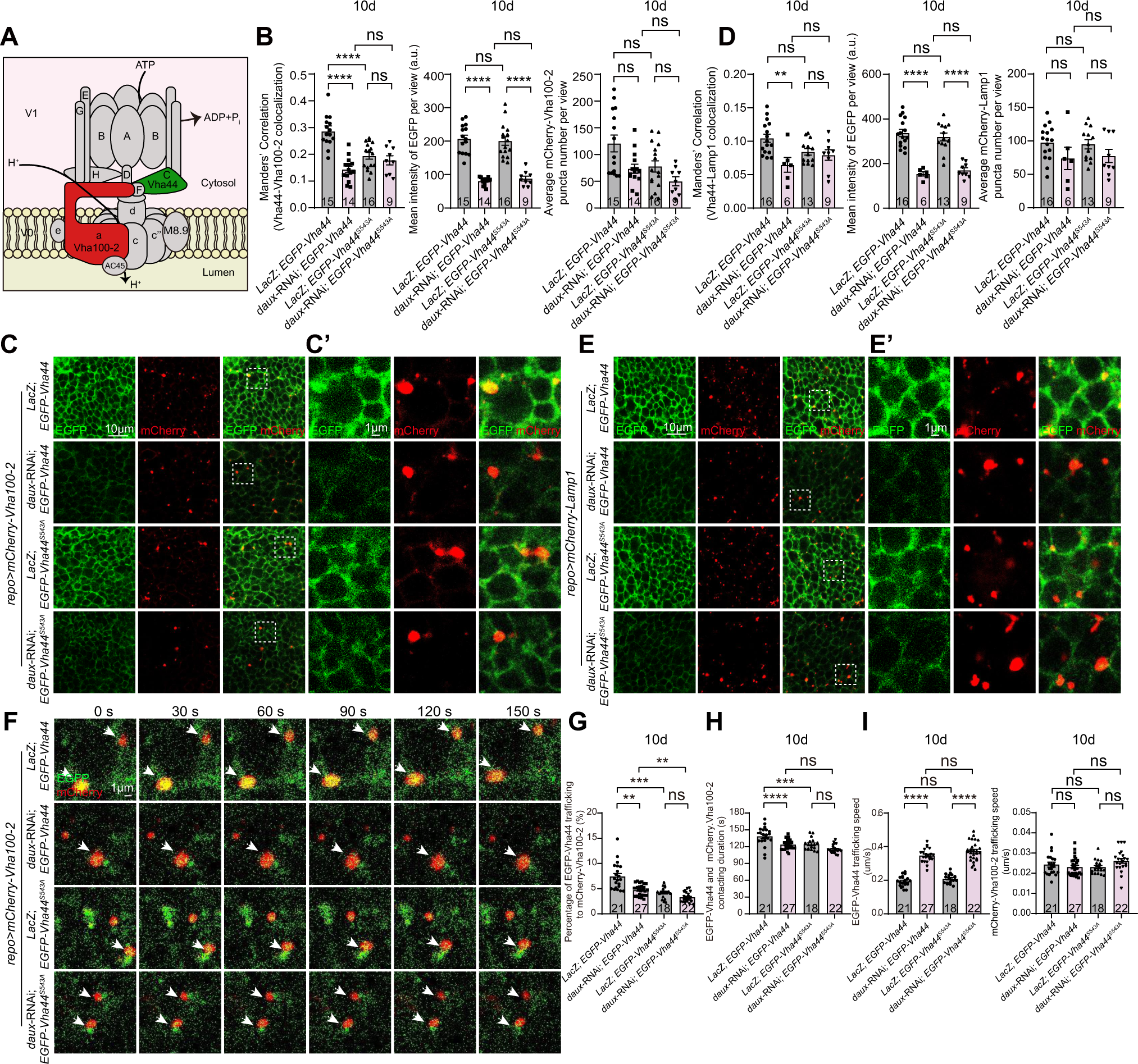

To analyze whether dAux-mediated Vha44 S543 phosphorylation affects the V-ATPase assembly, transgenic flies expressing the EGFP-Vha44^S543A^ variant carrying the alanine substitution of S543 were generated. Compared to EGFP-Vha44, EGFP-Vha44^S543A^ expression in glia reduced the Vha44^S543A^ colocalization with Vha100-2 or Lamp1, suggesting that non-phosphorylatable Vha44^S543A^ assemble with the V0 subcomplex less efficiently (Figures 5B-5E). Whereas *daux*-RNAi expression in glia reduced Vha44 colocalization with Vha100-2 or Lamp1, it failed to impact the Vha44^S543A^ colocalization with either (Figures 5B-5E). Live-cell imaging analysis also showed that compared to EGFP-Vha44, the percentage of EGFP-Vha44^S543A^ being trafficked to the vicinity of mCherry-Vha100-2 decreased (Figures 5F and 5G), and the duration of the Vha44^S543A^-Vha100-2 contact was shorter (Figures 5F and 5H). While lack of glial dAux increased the trafficking speed of EGFP-Vha44^S543A^-positive puncta (Figures 5F and 5I), the percentage of EGFP-Vha44^S543A^ being trafficked to mCherry-Vha100-2 or the duration of Vha44^S543A^-Vha100-2 contact remained unaffected (Figures 5F-5H). Taken together, these results suggest that Vha44 S543 phosphorylation underlies the mechanism of dAux-mediated V-ATPase assembly. It is noteworthy to mention that Manders’ Correlation was carefully chosen to analyze the colocalization of EGFP-Vha44 or EGFP-Vha44^S543A^ with Vha100-2 or Lamp1 as EGFP levels were affected by dAux (Figures 5B and 5D)^31–34^.

### dAux-mediated Vha44 S543 phosphorylation contributes to DA neurodegeneration and locomotor function

We previously reported that lack of glial dAux causes DA neurodegeneration at the PPM1/2 cluster, where DA neurons have been shown to degenerate upon α-syn overexpression^35^, and locomotor deficits, both pathological hallmarks implicated in PD^21^. Interestingly, whereas DA neurodegeneration at other clusters like protocerebral posterior lateral (PPL)1 and protocerebral posterior medial (PPM)3 was also detected^36, 37^, flies lacking glial dAux also exhibited shortened lifespan and sleep disturbance, one of the earlier PD-associated symptoms (Figures S8A-S8F). Consistent with flies lacking glial dAux, *Gak* cKO mice exhibited significant DA neuron loss in the substantia nigra (Figures S8G and S8H), locomotor deficits as quantified by the footprint analysis and pole test (Figures S8I and S8J), and body weight loss (Figure S8K). These results demonstrate that Gak/dAux function in glia is important for a broad spectrum of symptoms implicated in PD.

We then investigated if Vha44 and its S543 phosphorylation are also regulators of these cellular and behavioral symptoms. Interestingly, the number of DA neurons at the PPM1/2 cluster was significantly reduced in flies lacking Vha44 or expressing Myc-Vha44^S543A^ in glia in an age-dependent manner (Figures 6A-6E). Myc-Vha44 expression in glia caused an increase in the DA neuron number on day 3 but not on day 10 or 20 (Figures 6D and 6E). Furthermore, glial expression of Myc-Vha44, but not Myc-Vha44^S543A^, restored the *daux*-RNAi-induced DA neuron loss (Figures 6F and 6G). On the other hand, *vha44*-RNAi or Myc-Vha44^S543A^ expression in glia disrupted fly locomotion in an age-dependent manner, whereas Myc-Vha44 expression in glia affects fly locomotion at day 10 (Figures 6H and 6I). Re-introducing Myc-Vha44 expression in glia significantly rescued the *daux*-RNAi-induced locomotor deficits, whereas Myc-Vha44^S543A^ also did so but less potently (Figure 6J). Altogether, these results suggest that dAux-mediated Vha44 S543 phosphorylation underlies the mechanism of DA neurodegeneration and locomotor function implicated in PD.

**Figure 6.**
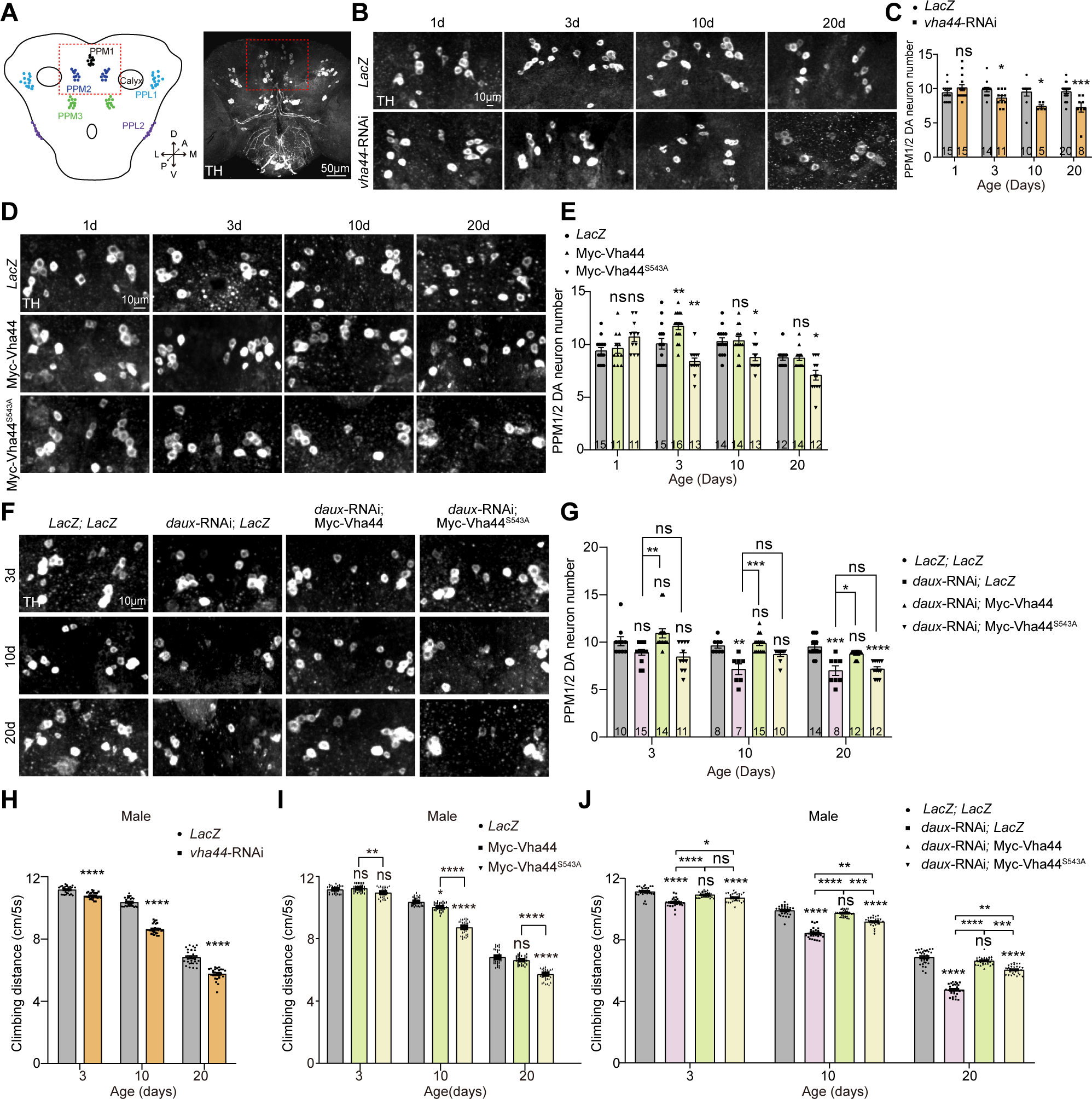

### Endogenous and PFF α-syn levels increase in *Gak* cKO microglia

Based on the results that α-syn levels increase in the absence of glial dAux (Figure 2) and that *Gak* cKO mice exhibit symptoms implicated in PD (Figure S8G-K), we further investigated the pathological relevance of glial Gak with PD by analyzing the α-syn levels in *Gak* cKO mice. Specifically, we used antibodies that detect the pathological component of the brain inclusions, the α-syn with phosphorylated serine 129 (pS129-α-syn), to quantitatively assess the α-syn levels^38–40^. Interestingly, whereas *Gak* cKO microglia exhibited shorter processes, higher CD68-positive phagocytic activity, and amoeboid-like structures in a potentially activated state (Figures 7A, C, D and E), pS129-α-syn levels and its colocalization with the microglial Iba1 were also elevated in *Gak* cKO microglia (Figures 7B and 7F). These accumulated pS129-α-syn were also highly colocalized with lysosomes in the *Gak* cKO microglia, reinforcing the notion that α-syn is degraded through a lysosome-dependent pathway in glia (Figures 7G and 7H).

**Figure 7.**
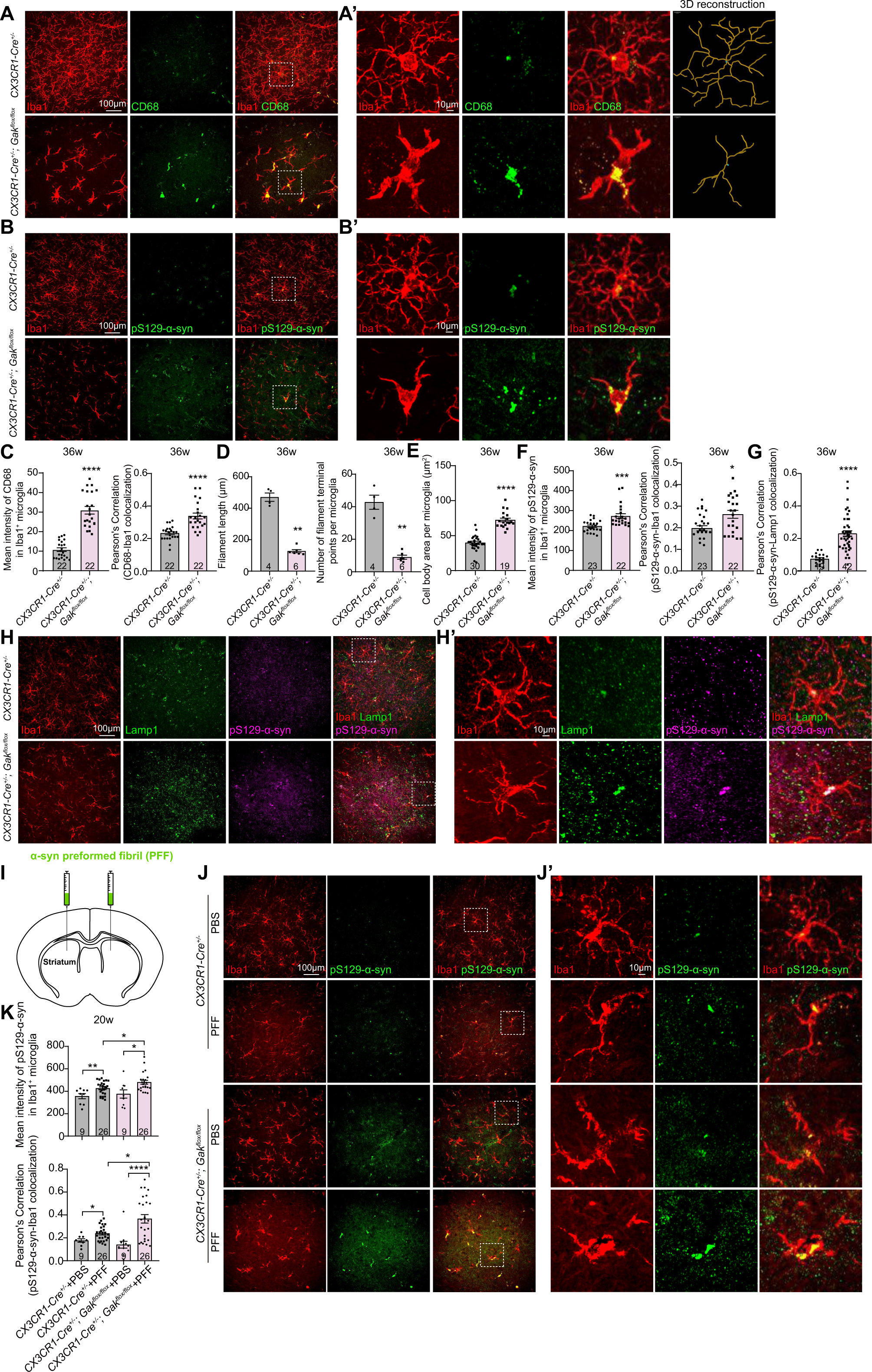

In addition, upon the bilateral inoculation of the α-syn preformed fibril (PFF) in the dorsal striatum, microglial pS129-α-syn intensities and the pS129-α-syn-Iba1 colocalization increased in the substantia nigra (+/− PFF), and further increased in the PFF-treated *Gak* cKO mice compared to the PFF-treated Cre alone controls (Figures 7I-7K). These results indicate that lacking microglial Gak promotes pS129-α-syn accumulation, reinforcing the notion that Gak regulates the degradation of pathological α-syn inclusions in glia.

## Discussion

Although neurodegeneration is the pathological hallmark for neurodegenerative diseases, glial responses constituting neuroimmune activation and signaling crosstalk during different stages have also widely contributed to the disease pathology. Furthermore, the intriguing phenomenon of protein inclusions observed in both neurons and glia raises substantial interest in how these pathological aggregates are formed, transmitted, and eliminated in different cell types. Our findings in flies and mice demonstrate that α-syn is degraded via a lysosomal pathway in glia; alternation in the lysosomal acidification/function impairs α-syn clearance. The disruption in lysosomal pH is caused by a phosphorylation-dependent regulation on the V-ATPase assembly, controlling the pumping of protons into the lysosomal lumen to construct the acidic milieu for substrate degradation. Mechanistically, lack of the PD risk factor dAux causes α-syn accumulation in glia, potentially owing to the lack of phosphorylation on the serine 543 of Vha44, the catalytic V1C subunit of the lysosomal V-ATPase. The V-ATPase assembly is therefore hindered, disrupting the lysosome pH for α-syn degradation (Figure 8).

**Figure 8.**
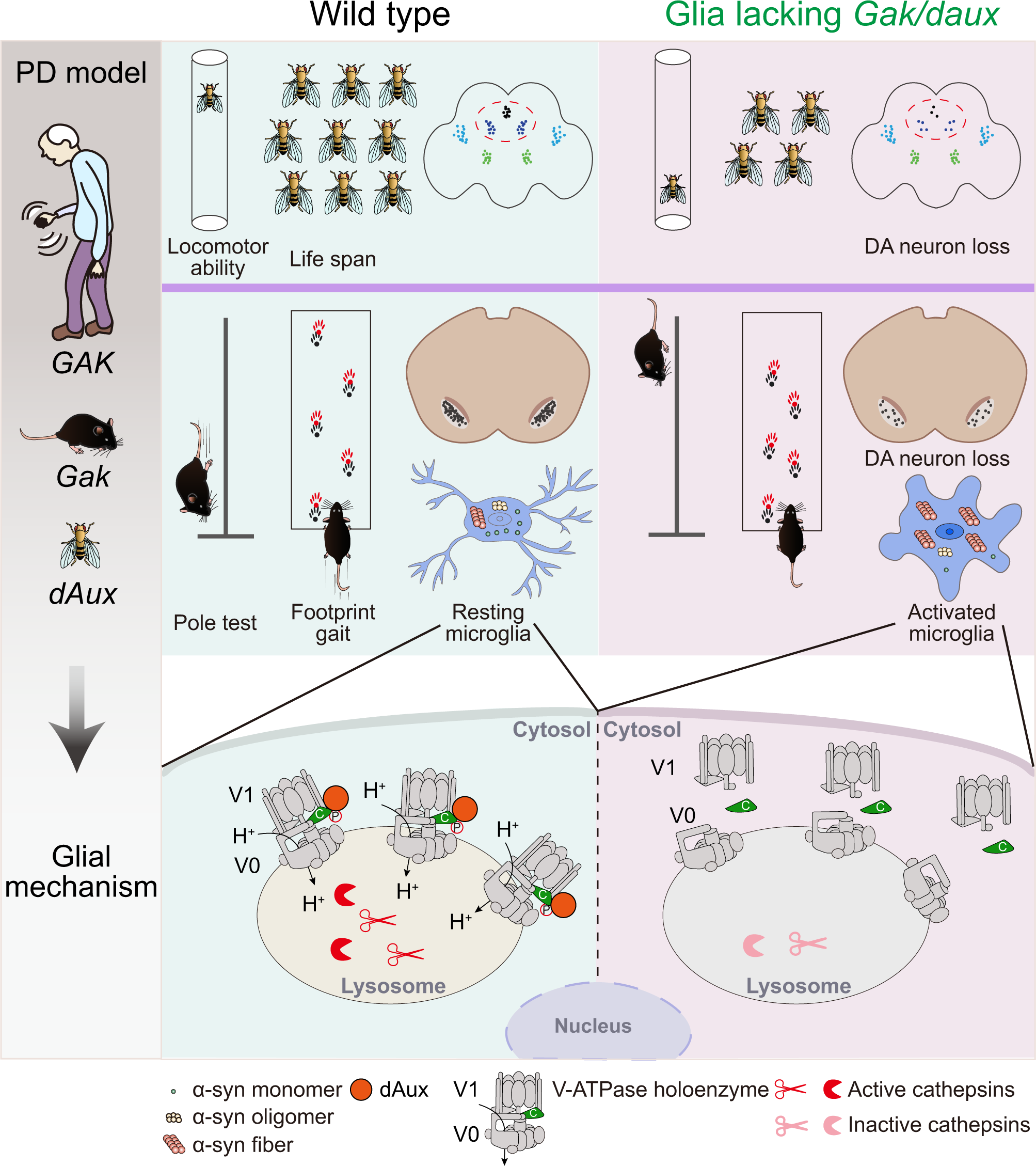

### V-ATPase subunit phosphorylation, assembly, and its pathological implication

The V-ATPase activity is modulated by the reversible association/dissociation of V1 and V0 subcomplexes in response to various conditions, such as glucose^41^, amino acid^42^, feeding^43^, serotonin (5-hydroxytryptamine, 5-HT)^44^, and cell maturation^45^. Previous reports have shown that some of the V-ATPase subunits are targets for phosphorylation. For instance, AP2 interacts and phosphorylates the V1B subunit^46^; PKA or AMP-activated protein kinase (AMPK) phosphorylates the V1A subunit^47, 48^; *Arabidopsis* AtWNK8 phosphorylates the V1C subunit at multiple sites^49^. Nonetheless, how these modifications on different V-ATPase subunits impact on its activity and function remains largely unknown. While some of the subunits are phosphorylated in isolation, some might require to be part of the complex allowing phosphorylation occurs^48, 50^. Our present study uncovers a phosphorylation-dependent switch of the V-ATPase assembly, linking the subunit phosphorylation with the activity of the whole enzyme.

Dysregulation of V-ATPase activity has been shown in many neurodegenerative disorders including PD^19, 51^. For instance, the modification, protein level, or protein-protein interaction of the V0a1 subunit is critical for the pathology of different types of neurodegenerative diseases^52–55^. The PD risk factor leucine-rich repeat kinase 2 (LRRK2) also regulates V0a1 protein level and localization^53^. Interestingly, our current findings reveal a critical role of the V1C subunit. Lack of Vha44 or blocking Vha44 S543 phosphorylation causes impaired lysosomal acidification and hydrolase activity, constructing unfavorable venues for α-syn degradation. Furthermore, flies exhibit DA neurodegeneration and locomotor deficits in the absence of Vha44. These findings indicate that a wrongly modified V-ATPase subunit hinders its proper assembly and impairs lysosomal acidification, leading to potential α-syn accumulation and ultimately PD-like symptoms. Of note, GAK interacts with LRRK2^56^, yet these two proteins modulate the V-ATPase activity via different subunits: LRRK2 interacts with V0a1 whereas dAux interacts with V1C, potentiating different V-ATPase subunit-mediated mechanisms underlying PD.

### Gak/dAux functions as a kinase regulating lysosomal function in glia

We identified the *Drosophila* GAK homolog, dAux, functioning as a kinase in phosphorylating Vha44 at S543. In the absence of dAux, Vha44 S543 phosphorylation is significantly reduced. Interestingly, glial expression of Vha44, but not the non-phosphorylatable Vha44 variant Vha44^S543A^, in adult fly brains lacking glial dAux, re-acidify lysosomes, pointing to a critical role of Vha44 S543 phosphorylation in dAux-mediated lysosomal acidification. On the other hand, glial expression of the dAux lacking the kinase domain fails to rescue the defects in lysosomal acidification in the absence of dAux, further reinforcing the requirement of dAux kinase function in regulating the glial lysosomal pathway.

Recently, we reported a role for dAux in autophagy in glia^21^. These findings reveal that dAux regulates the initiation of autophagy by interacting with the master regulator ULK1/Atg1. Despite the physical interaction, we failed to detect significant changes in Atg1 phosphorylation in the absence of glial dAux, suggesting that dAux regulates ULK1/Atg1 activity in a kinase-independent manner. Thus, our previous and present studies uncover two independent mechanisms mediated by dAux, regulating different steps along the glial degradation pathway.

### A glia-specific lysosomal pathway associated with α-syn degradation in PD

Targeting protein aggregates and inclusions in the degenerative brains has been considered as a strategic means for ameliorating disease progression. Our findings uncover the mechanism of a potential PD risk factor regulating α-syn clearance in glia. Considering the V-ATPase disassembly, lysosomal defects, and brain α-syn accumulation (prominently seen in lysosomes) observed in the absence of glial dAux, the mechanism of dAux-mediated Vha44 S543 phosphorylation likely constitutes a lysosomal pathway associated with α-syn clearance in glia.

Despite multiple SNPs have been identified in the locus containing *GAK*, implicating its clinical relevance^57–59^, whether *GAK* is a PD risk gene remains controversial. Pathologically, potential *GAK* mutations could happen in both neurons and glia in PD. Yet, our observations on how the lysosomal acidification remains largely intact upon neuronal dAux inhibition suggest the glia specificity of the present mechanism (Figure S4). Given that GAK/dAux is a multifunctional protein, it is possible that GAK/dAux functions differently in neurons and controls downstream pathways other than lysosome. As GAK/dAux function in neurons remains largely unclear, we speculate that GAK/dAux function in neurons and glia are separate and independent from each other, whereas dAux-mediated lysosomal pathway contributes to α-syn degradation in glia.

While most of the hypotheses underlying PD pathology are often driven by a neuronal origin of dysfunctional machineries or cellular defects, glia have only recently received more attention as possible drivers for neurodegeneration^7^. For instance, the AD risk factor gene *Apolipoprotein E* (*APOE*) is predominantly expressed in astrocytes^60^ and its genetic mutations are associated with an increasing risk of developing AD, supporting a role for genetic causes from glia driving neurodegeneration. Many PD risk genes, like *LRRK2*, *atg9*, and *vps13*, are also expressed in glia, and modulate neurodegeneration and α-syn accumulation in a non-cell-autonomous manner^61–64^. Our findings implicate another potential PD risk factor functioning in a glial context. Lack of glial Gak/dAux leads to a broad spectrum of symptoms implicated in PD in both flies and mice, including DA neurodegeneration, locomotor dysfunction, shortened lifespan, microglial activation, and some of the earlier symptoms such as sleep defects. Consistently, DA neurodegeneration and locomotor deficits are also detected in the absence of glial Vha44. Glial expression of Vha44, but not the non-phosphorylatable Vha44 variant, restores these deficits. In combination with our cellular observations, these behavioral deficits together demonstrate a kinase-dependent lysosomal pathway in glia regulating the symptoms implicated in PD.

The present study identifies a glia-specific pathway for the lysosomal degradation of α-syn. Our findings elucidate the underlying mechanism of PD by uncovering a cell-type specific clearance program for the brain α-syn inclusions. With accumulating evidence demonstrating the importance of glia in neurodegenerative diseases, these findings help to pinpoint potential therapeutical targets from a glial perspective to ameliorate PD.

## Acknowledgements

We thank Bloomington *Drosophila* Stock Center, Vienna *Drosophila* RNAi Center, the Core Facility of *Drosophila* Resource and Technology, Shanghai Institute of Biochemistry and Cell Biology, Chinese Academy of Sciences, Yufeng Pan, and Aike Guo for fly stocks; Chih-Hao Lee for IMG cell line; Jiawei Zhou for *CX3CR1-Cre* mice. We also thank the Molecular Imaging Core Facility (MICF), the Molecular and Cell Biology Core Facility (MCBCF), and the Multi-Omics Core Facility (MOCF) at the School of Life Science and Technology, ShanghaiTech University for providing technical support; Cong Liu for PFF synthesis; Yu Kong and Lijun Pan in Electron Microscopy Facilities of Center for Excellence in Brain Science and Technology, Chinese Academy of Science for assistance with TEM sample preparation; DroBot Biotechnology for quality fly food supply, delicate fly-keeping service, and experimental device design; Ho lab members for discussion and comments. This work was supported by grants from ShanghaiTech and National Yang Ming Chiao Tung University (intramural), National Natural Science Foundation of China (32170962, 32271004, 32371063, and 32071009), 2030 Cross-Generation International Outstanding Young Scholars Program National Science and Technology Council Taiwan (113-2628-B-A49-007), and Brain Research Center National Yang Ming Chiao Tung University from The Featured Areas Research Center Program within the framework of the Higher Education Sprout Project by the Ministry of Education (MOE) in Taiwan.

## Author contributions

S.Z, L.W, S.Y, and M.S.H conceived and designed the study. S.Z, L.W, S.Y, H.W, S.L, R.W, Y.L, and W.Y performed the experiments. S.Z, L.W, S.Y, Y.T, C.L, K.W.H, and M.S.H analyzed the data. S.Z, L.W, S.Y, and M.S.H wrote the paper. All authors read and approved the manuscript.

## Data Availability Statement

All the data supporting this study are available from the corresponding author upon reasonable request.

## Declaration of Interests

The authors declare no competing interests.

## Notes

### Competing Interest Statement

The authors have declared no competing interest.

### Summary of Updates

We have updated some results and re-organized figures

## References

1. Jellinger KA. Multiple System Atrophy: An Oligodendroglioneural Synucleinopathy1. J Alzheimers Dis 2018; 62:1141–79.

2. Peng C, Trojanowski JQ, Lee VM. Protein transmission in neurodegenerative disease. Nat Rev Neurol 2020; 16:199–212.

3. Valdinocci D, Radford RA, Siow SM, Chung RS, Pountney DL. Potential Modes of Intercellular alpha-Synuclein Transmission. Int J Mol Sci 2017; 18.

4. Tanriover G, Bacioglu M, Schweighauser M, Mahler J, Wegenast-Braun BM, Skodras A, et al. Prominent microglial inclusions in transgenic mouse models of alpha-synucleinopathy that are distinct from neuronal lesions. Acta Neuropathol Commun 2020; 8:133.

5. Ho MS. Microglia in Parkinson’s Disease. Adv Exp Med Biol 2019; 1175:335–53.

6. Subhramanyam CS, Wang C, Hu Q, Dheen ST. Microglia-mediated neuroinflammation in neurodegenerative diseases. Semin Cell Dev Biol 2019; 94:112–20.

7. Gleichman AJ, Carmichael ST. Glia in neurodegeneration: Drivers of disease or along for the ride? Neurobiol Dis 2020; 142:104957.

8. Cheng J, Lu Q, Song L, Ho MS. alpha-Synuclein Trafficking in Parkinson’s Disease: Insights From Fly and Mouse Models. ASN Neuro 2018; 10:1759091418812587.

9. Filippini A, Gennarelli M, Russo I. alpha-Synuclein and Glia in Parkinson’s Disease: A Beneficial or a Detrimental Duet for the Endo-Lysosomal System? Cell Mol Neurobiol 2019; 39:161–8.

10. Yi S, Wang L, Wang H, Ho MS, Zhang S. Pathogenesis of alpha-Synuclein in Parkinson’s Disease: From a Neuron-Glia Crosstalk Perspective. Int J Mol Sci 2022; 23.

11. Ballabio A, Bonifacino JS. Lysosomes as dynamic regulators of cell and organismal homeostasis. Nat Rev Mol Cell Biol 2020; 21:101–18.

12. Paudel RR, Lu D, Roy Chowdhury S, Monroy EY, Wang J. Targeted Protein Degradation via Lysosomes. Biochemistry 2023; 62:564–79.

13. Yang C, Wang X. Lysosome biogenesis: Regulation and functions. J Cell Biol 2021; 220.

14. Nishi T, Forgac M. The vacuolar (H+)-ATPases--nature’s most versatile proton pumps. Nat Rev Mol Cell Biol 2002; 3:94–103.

15. Beyenbach KW, Wieczorek H. The V-type H+ ATPase: molecular structure and function, physiological roles and regulation. J Exp Biol 2006; 209:577–89.

16. Stevens TH, Forgac M. Structure, function and regulation of the vacuolar (H+)-ATPase. Annu Rev Cell Dev Biol 1997; 13:779–808.

17. Futai M, Sun-Wada GH, Wada Y, Matsumoto N, Nakanishi-Matsui M. Vacuolar-type ATPase: A proton pump to lysosomal trafficking. Proc Jpn Acad Ser B Phys Biol Sci 2019; 95:261–77.

18. Cotter K, Stransky L, McGuire C, Forgac M. Recent Insights into the Structure, Regulation, and Function of the V-ATPases. Trends Biochem Sci 2015; 40:611–22.

19. Song Q, Meng B, Xu H, Mao Z. The emerging roles of vacuolar-type ATPase-dependent Lysosomal acidification in neurodegenerative diseases. Transl Neurodegener 2020; 9:17.

20. Forgac M. Vacuolar ATPases: rotary proton pumps in physiology and pathophysiology. Nat Rev Mol Cell Biol 2007; 8:917–29.

21. Zhang S, Yi S, Wang L, Li S, Wang H, Song L, et al. Cyclin-G-associated kinase GAK/dAux regulates autophagy initiation via ULK1/Atg1 in glia. Proc Natl Acad Sci U S A 2023; 120:e2301002120.

22. Wang L, Zhang S, Yi S, Ho MS. A new regulator of autophagy initiation in glia. Autophagy 2023:1–3.

23. Miyazaki M, Hiramoto M, Takano N, Kokuba H, Takemura J, Tokuhisa M, et al. Targeted disruption of GAK stagnates autophagic flux by disturbing lysosomal dynamics. Int J Mol Med 2021; 48.

24. Munson MJ, Mathai BJ, Ng MYW, Trachsel-Moncho L, de la Ballina LR, Schultz SW, et al. GAK and PRKCD are positive regulators of PRKN-independent mitophagy. Nat Commun 2021; 12:6101.

25. Senturk M, Lin G, Zuo Z, Mao D, Watson E, Mikos AG, Bellen HJ. Ubiquilins regulate autophagic flux through mTOR signalling and lysosomal acidification. Nat Cell Biol 2019; 21:384–96.

26. Kovacs AL, Reith A, Seglen PO. Accumulation of autophagosomes after inhibition of hepatocytic protein degradation by vinblastine, leupeptin or a lysosomotropic amine. Exp Cell Res 1982; 137:191–201.

27. Mauvezin C, Nagy P, Juhasz G, Neufeld TP. Autophagosome-lysosome fusion is independent of V-ATPase-mediated acidification. Nat Commun 2015; 6:7007.

28. Zhang Y, Chen K, Sloan SA, Bennett ML, Scholze AR, O’Keeffe S, et al. An RNA-sequencing transcriptome and splicing database of glia, neurons, and vascular cells of the cerebral cortex. J Neurosci 2014; 34:11929–47.

29. Zhang Y, Sloan SA, Clarke LE, Caneda C, Plaza CA, Blumenthal PD, et al. Purification and Characterization of Progenitor and Mature Human Astrocytes Reveals Transcriptional and Functional Differences with Mouse. Neuron 2016; 89:37–53.

30. McCarthy RC, Lu DY, Alkhateeb A, Gardeck AM, Lee CH, Wessling-Resnick M. Characterization of a novel adult murine immortalized microglial cell line and its activation by amyloid-beta. J Neuroinflammation 2016; 13:21.

31. Bido S, Muggeo S, Massimino L, Marzi MJ, Giannelli SG, Melacini E, et al. Microglia-specific overexpression of alpha-synuclein leads to severe dopaminergic neurodegeneration by phagocytic exhaustion and oxidative toxicity. Nat Commun 2021; 12:6237.

32. Nabar NR, Heijjer CN, Shi CS, Hwang IY, Ganesan S, Karlsson MCI, Kehrl JH. LRRK2 is required for CD38-mediated NAADP-Ca(2+) signaling and the downstream activation of TFEB (transcription factor EB) in immune cells. Autophagy 2022; 18:204–22.

33. Winslow AR, Chen CW, Corrochano S, Acevedo-Arozena A, Gordon DE, Peden AA, et al. alpha-Synuclein impairs macroautophagy: implications for Parkinson’s disease. J Cell Biol 2010; 190:1023–37.

34. Rivera OC, Hennigar SR, Kelleher SL. ZnT2 is critical for lysosome acidification and biogenesis during mammary gland involution. Am J Physiol Regul Integr Comp Physiol 2018; 315:R323–R35.

35. Feany MB, Bender WW. A Drosophila model of Parkinson’s disease. Nature 2000; 404:394–8.

36. Budnik V, White K. Catecholamine-containing neurons in Drosophila melanogaster: distribution and development. J Comp Neurol 1988; 268:400–13.

37. Nässel DR, Elekes K. Aminergic neurons in the brain of blowflies and Drosophila: dopamine- and tyrosine hydroxylase-immunoreactive neurons and their relationship with putative histaminergic neurons. Cell Tissue Res 1992; 267:147–67.

38. Fujiwara H, Hasegawa M, Dohmae N, Kawashima A, Masliah E, Goldberg MS, et al. alpha-Synuclein is phosphorylated in synucleinopathy lesions. Nat Cell Biol 2002; 4:160–4.

39. Smith WW, Margolis RL, Li X, Troncoso JC, Lee MK, Dawson VL, et al. α-Synuclein Phosphorylation Enhances Eosinophilic Cytoplasmic Inclusion Formation in SH-SY5Y Cells. The Journal of Neuroscience 2005; 25:5544–52.

40. Walker DG, Lue L-F, Adler CH, Shill HA, Caviness JN, Sabbagh MN, et al. Changes in properties of serine 129 phosphorylated α-synuclein with progression of Lewy-type histopathology in human brains. Experimental Neurology 2013; 240:190–204.

41. Kane PM. Disassembly and reassembly of the yeast vacuolar H(+)-ATPase in vivo. J Biol Chem 1995; 270:17025–32.

42. Stransky LA, Forgac M. Amino Acid Availability Modulates Vacuolar H+-ATPase Assembly. J Biol Chem 2015; 290:27360–9.

43. Sumner JP, Dow JA, Earley FG, Klein U, Jager D, Wieczorek H. Regulation of plasma membrane V-ATPase activity by dissociation of peripheral subunits. J Biol Chem 1995; 270:5649–53.

44. Zimmermann B, Dames P, Walz B, Baumann O. Distribution and serotonin-induced activation of vacuolar-type H+-ATPase in the salivary glands of the blowfly Calliphora vicina. J Exp Biol 2003; 206:1867–76.

45. Trombetta ES, Ebersold M, Garrett W, Pypaert M, Mellman I. Activation of lysosomal function during dendritic cell maturation. Science 2003; 299:1400–3.

46. Myers M, Forgac M. The coated vesicle vacuolar (H+)-ATPase associates with and is phosphorylated by the 50-kDa polypeptide of the clathrin assembly protein AP-2. J Biol Chem 1993; 268:9184–6.

47. Alzamora R, Thali RF, Gong F, Smolak C, Li H, Baty CJ, et al. PKA regulates vacuolar H+-ATPase localization and activity via direct phosphorylation of the a subunit in kidney cells. J Biol Chem 2010; 285:24676–85.

48. Alzamora R, Al-Bataineh MM, Liu W, Gong F, Li H, Thali RF, et al. AMP-activated protein kinase regulates the vacuolar H+-ATPase via direct phosphorylation of the A subunit (ATP6V1A) in the kidney. Am J Physiol Renal Physiol 2013; 305:F943–56.

49. Hong-Hermesdorf A, Brux A, Gruber A, Gruber G, Schumacher K. A WNK kinase binds and phosphorylates V-ATPase subunit C. FEBS Lett 2006; 580:932–9.

50. Voss M, Vitavska O, Walz B, Wieczorek H, Baumann O. Stimulus-induced phosphorylation of vacuolar H(+)-ATPase by protein kinase A. J Biol Chem 2007; 282:33735–42.

51. Colacurcio DJ, Nixon RA. Disorders of lysosomal acidification-The emerging role of v-ATPase in aging and neurodegenerative disease. Ageing Res Rev 2016; 32:75–88.

52. Avrahami L, Farfara D, Shaham-Kol M, Vassar R, Frenkel D, Eldar-Finkelman H. Inhibition of glycogen synthase kinase-3 ameliorates beta-amyloid pathology and restores lysosomal acidification and mammalian target of rapamycin activity in the Alzheimer disease mouse model: in vivo and in vitro studies. J Biol Chem 2013; 288:1295–306.

53. Wallings R, Connor-Robson N, Wade-Martins R. LRRK2 interacts with the vacuolar-type H+-ATPase pump a1 subunit to regulate lysosomal function. Hum Mol Genet 2019; 28:2696–710.

54. Lee JH, Yu WH, Kumar A, Lee S, Mohan PS, Peterhoff CM, et al. Lysosomal proteolysis and autophagy require presenilin 1 and are disrupted by Alzheimer-related PS1 mutations. Cell 2010; 141:1146–58.

55. Im E, Jiang Y, Stavrides PH, Darji S, Erdjument-Bromage H, Neubert TA, et al. Lysosomal dysfunction in Down syndrome and Alzheimer mouse models is caused by v-ATPase inhibition by Tyr(682)-phosphorylated APP betaCTF. Sci Adv 2023; 9:eadg1925.

56. Beilina A, Rudenko IN, Kaganovich A, Civiero L, Chau H, Kalia SK, et al. Unbiased screen for interactors of leucine-rich repeat kinase 2 supports a common pathway for sporadic and familial Parkinson disease. Proc Natl Acad Sci U S A 2014; 111:2626–31.

57. Lin CH, Chen ML, Tai YC, Yu CY, Wu RM. Reaffirmation of GAK, but not HLA-DRA, as a Parkinson’s disease susceptibility gene in a Taiwanese population. Am J Med Genet B Neuropsychiatr Genet 2013; 162B:841–6.

58. Rhodes SL, Sinsheimer JS, Bordelon Y, Bronstein JM, Ritz B. Replication of GWAS associations for GAK and MAPT in Parkinson’s disease. Ann Hum Genet 2011; 75:195–200.

59. Tseng WE, Chen CM, Chen YC, Yi Z, Tan EK, Wu YR. Genetic variations of GAK in two Chinese Parkinson’s disease populations: a case-control study. PLoS One 2013; 8:e67506.

60. Xu Q, Bernardo A, Walker D, Kanegawa T, Mahley RW, Huang Y. Profile and regulation of apolipoprotein E (ApoE) expression in the CNS in mice with targeting of green fluorescent protein gene to the ApoE locus. J Neurosci 2006; 26:4985–94.

61. Wang L, Wang H, Yi S, Zhang S, Ho MS. A LRRK2/dLRRK-mediated lysosomal pathway that contributes to glial cell death and DA neuron survival. Traffic 2022; 23:506–20.

62. Yi S, Wang L, Ho MS, Zhang S. The autophagy protein Atg9 functions in glia and contributes to parkinsonian symptoms in a Drosophila model of Parkinson’s disease. Neural Regen Res 2024; 19:1150–5.

63. Olsen AL, Feany MB. Glial alpha-synuclein promotes neurodegeneration characterized by a distinct transcriptional program in vivo. Glia 2019; 67:1933–57.

64. Olsen AL, Feany MB. Parkinson’s disease risk genes act in glia to control neuronal alpha-synuclein toxicity. Neurobiol Dis 2021; 159:105482.

