## Supplementary Materials for "dAux orchestrates the phosphorylation-dependent assembly of the lysosomal V-ATPase in glia and contributes to α-synuclein degradation"

Shiping Zhang^1,4,†^, Linfang Wang^1,†^, Shuanglong Yi^1,5†^, Yu-ting Tsai^2,3^, Honglei Wang^1^, Shuhua Li^1^, Ruiqi Wang^1,6^, Yang Liu^7^, Wei Yan^8^, Chang Liu^8^, Kai-Wen He^7^, Margaret S. Ho^1,2,3,*^

**This docx file includes:**

Materials and Methods

Supplementary Figures S1-S8

Table S1

**Materials and Methods**

***Drosophila* genetics**

Flies were raised under standard yeast-cornmeal-agar medium at 25℃ and 70% humidity. All fly crosses were carried out at 25°C with standard laboratory conditions. Fly strains used include: *w^1118^* (BL#5905)*, GMR57C10-*lexA (BL#81080), *UAS-LacZ* (BL#1777), *UAS-luc-*RNAi (BL#31603)*, UAS-daux-*RNAi (V#16182), *UAS-daux-*RNAi^#2^ (BL#39017), *UAS-vha44-*RNAi (BL#33884), *UAS-HRP* (gift from Aike Guo), *repo-*GAL4^1^, and *UAS-Lamp1-GFP*^2^. Detailed fly genotypes used in this study were listed in Table S1.

**Mouse genetics**

The B6;129S6-*Gak^tm2Legr^*/Mmjax mice (Cat. #36793-JAX, the Jackson Laboratory, Bar Harbor, ME, USA) have loxP sites flanking *cyclin G associated kinase* (*Gak*) gene exon 1. *CX3CR1-Cre* transgenic mice were gifts from Jiawei Zhou. *CX3CR1-Cre^+/–^* littermates were used as controls. All mice were maintained with unrestricted access to food and water under a 12 hours light/dark cycle at 22 ± 2°C. All experimental protocols were approved and directed by the Animal Care and Use Committee of National Center for Protein Science Shanghai.

**Cell culture and transfections**

IMG cells: IMG cell line was a gift from Chih-Hao Lee^3^. Cells were cultured with high glucose (4.5 g/L), 10% fetal bovine serum (Cat. #10099141, Gibco) and 100 units/mL penicillin-streptomycin (Cat. #15140-122, Gibco) in Dulbecco’s modified Eagle medium (Cat. #11965092, Gibco, New York, NY, USA) at 37°C with 5% CO2 humidified atmosphere. IMG cells were transfected with Lipo 2000 (Cat. #11668019, Life Technologies, Carlsbad, CA, USA) according to the manufacturer’s protocol. Briefly, a 12-well plate was used to seed IMG cells overnight, and then incubated with the mixed 40 pmol siRNA (GenePharma) and 2 ul Lipo 2000 which was diluted and reacted at room temperature for 20 minutes in 100 µl Opti-MEM (Cat. #31985070, Gibco). After 6 hours culture at 37°C with 5% CO2, cells were changed into growth medium and harvested after 72 hours.

*Drosophila* S2 cells: S2 cells were cultured in Schneider’s medium (Cat. #21720024, Gibco) at 28°C and the Effectene Transfection Reagent (Cat. #301425, Qiagen) was used for transfection. After seeding for 6 hours, cells were transfected, 24 hours later, CuSO₄ were added at a final concentration of 1mM, and cells were harvested at another 24 hours.

**Molecular biology**

Plasmid cloning: Plasmids were generated by Polymerase chain reaction (PCR) or gene synthesis by Genscript (Nanjing, China). The plasmids expressing Vha44, Vha44^S543A^, Vha100-2, Lamp1, or human α-syn were subcloned into 6xMyc-, EGFP-, or mCherry- tagged *pUAST-attB* vector, or 6xMyc-tagged *pJFRC19-13xlexAop2-attB* vector (Cat. #26224, Addgene, Watertown, MA, USA). Fly microinjection was performed by the *Drosophila* Core Facility, Institute of Biochemistry and Cell Biology, Chinese Academy of Sciences.

Primers used for amplification:

*SNCA*-F: CCGGGCCTGGGATGTATTCATGAAAGGACTTTCAAAGG

*SNCA*-R: ACAAAGATCCTCTAGATTAGGCTTCAGG

qRT-PCR: Total RNAs of *Drosophila* heads were extracted with TransZol Up (Cat. #ET111-01, TransGen, Beijing, China). For reverse transcription, HiScript III RT SuperMix (Cat. #R323-01, Vazyme, Nanjing, China) kit was used. qRT-PCR reactions were conducted by ChamQ Universal SYBR qPCR Master Mix (Cat. #Q711-02, Vazyme) and ABI 7500 RT-PCR system. *rp49* was used as internal control. The expression levels were quantified by ΔΔCT method.

Primers used are listed below:

Flies:

*rp49*-F: CCACCAGTCGGATCGATATGC

*rp49*-R: CTCTTGAGAACGCAGGCGACC

*vha44*-F: TTGGTTCGTTGGCTGAAGGT

*vha44*-R: GCACGGATTCCACAAACACG

Antibody generation: The anti-pVha44^S543^ antibodies were produced by ABclonal Biotechnology co., Ltd (Wuhan, China) using a modified peptides EFPL(S-p)PAEL corresponding to the 539-547 amino acids of Vha44 isoform F, the peptides were injected into rabbits to generate the antibodies.

**Immunohistochemistry**

Adult fly brains: Adult fly brains at 10 days after eclosion were dissected and fixed for 40 minutes in 4% formaldehyde, washed for 3 times using PBT (PBS + 0.1% TX-100) and dissected further to devoid any debris. Then brains were blocked with 5% Normal Donkey Serum (NDS) in PBT solution and subsequently incubated overnight with primary antibodies at 4℃ and then secondary antibodies at room temperature for 2 hours. Primary antibodies used in the following concentrations: rabbit anti-Myc (1:2000, Cat. #0912-2, Hua An Biotechnology), and rabbit anti-tyrosine hydroxylase (1:300, Cat. #AB152, Merck), rabbit anti-Ref(2)P (1:500, Cat. #ab178440, Abcam, Cambridge, UK), and mouse anti-mono- and polyubiquitinylated conjugates antibody (FK2, 1:500, Cat. #BML-PW8810, Enzo Life Sciences, Farmingdale, NY, USA).

IMG cells: Glass coverslips in 12-well plates were used to seed cells the day before fixation with cold methanol (Cat. #100141190, Sinopharm, Beijing, China) for 15 minutes, followed by washing for 15 minutes with PBT (PBS + 0.1% TX-100), then blocked for 1 hour in PBS with 5% NDS. Cells were stained overnight with primary antibody at 4℃ and then secondary antibodies for 2 hours at room temperature. Primary antibodies were used in the following concentrations: rabbit anti-Lamp1 (1:400, Cat. #ab24170, Abcam). Nuclei were labeled by DAPI (Cat. #C1005, Beyotime, Shanghai, China) at room temperature for 5-10 minutes.

Mouse brain slices: Whole brain was collected following transcardial perfusion with isoflurane and perfused with 0.9% saline, followed by 4% paraformaldehyde in PBS. Next, whole brain tissue was post-fixed in 4% paraformaldehyde for 24 hours and then immersed in 30% (w/v) sucrose in PBS until the tissue settled to the bottom of the tube (~ 48 hours). A series of coronal sections (40 μm thick) across the Substantia Nigra were cut by a Thermo CryoStar NX50 cryostat (Thermofisher Scientific, USA). The sections were permeabilized with 0.3% Triton X-100 in PBS containing 5% normal goat serum at room temperature for 15 minutes. Sections were blocked with 5% normal goat serum in PBS and were then incubated with primary antibodies at 4°C overnight. The following day, samples were incubated with secondary antibodies conjugated with Alexa-fluorescein for 1 hour. Primary antibodies were used in the following concentrations: rabbit anti-TH (1:500, Cat. #AB152, Merck), rabbit anti-Iba1 (1:500, Cat. #019-19741, Wako), chicken anti-Iba1 (1:500, Cat. #234009, Synaptic System), mouse anti-CD68 (1:250, Cat. #2342153, Invitrogen, Carlsbad, CA, USA), rat anti-Lamp1 (1:200, Cat. #1D4B, DSHB), and rabbit anti-pS129-α-synuclein (1:250, Cat. #ab51253, Abcam).

For all samples, secondary antibodies used were from Jackson ImmunoResearch (West Grove, PA, USA): donkey anti-mouse Cy3 (1:1000, Cat. #715-166-151), donkey anti-rabbit Cy3 (1:1000, Cat. #711-166-152), donkey anti-rat Cy3 (1:1000, Cat. #712-165-153), goat anti-rabbit FITC (1:1000, Cat. #111-095-003), goat anti-chicken FITC (1:500, Cat. #103-005-155), and donkey anti-rat Cy5 (1:500, Cat. #712-175-150).

**LysoTracker staining**

After dissection in PBS, live adult fly brains were incubated in LysoTracker Green DND-26 (Cat. #L7526, Life technologies) or LysoTracker Red DND-99 (Cat. # 40739ES50, Yeasen, Shanghai, China) solution for 5 minutes at 25°C. The LysoTracker dyes were reconstituted with DMSO to form the stock solution, and then diluted 1:1000 with PBS for the final working concentration prior to use. Images were obtained immediately after the staining.

**Cathepsin B detection with Magic Red**

After dissection in PBS, live adult fly brains were incubated in Magic Red Cathepsin B (Magic Red® Cathepsin B assay kits, Cat. #ICT937, BioRad) solution for 5 minutes at 25°C. The Magic Red Cathepsin B was reconstituted with DMSO to form the stock solution, and then diluted 1:10 with diH2O for the final working concentration prior to use. Images were obtained immediately after the staining.

***In-vivo* live cell imaging**

Live adult fly brains were dissected and held steady by Vaseline in PBS, the puncta were immediately recorded in a single focal plane for 5 minutes using Nikon C2 microscope (40x water objective NA＝0.8w). Imaris (Bitplane, Zurich, Switzerland) was used to analyzed puncta trafficking speed and percentage with an Autoregressive Motion mode. The time for contacting durations were analyzed manually.

**Transmission electron microscopy**

Dissected adult fly brains were fixed, embedded, stained, and dehydrated as described^4^. Each brain was cut at 70 nm thick with a Diatomediamond knife on a Leica EM UC7 ultra-microtome (Leica Microsystems, Germany) and collected on copper grids. Images were taken by Talos L120C transmission electron microscope (Thermo Fisher Scientific, Waltham, MA, UAS) through an acceleration voltage of 120 kV.

**Biochemistry**

Western blot analysis: adult fly heads were homogenized by a motorized pestle (Cat. #116005500, MP Biomedicals, Irvine, CA, USA) in lysis buffer (0.4% NP-40, 0.2 mM EDTA, 150 mM NaCl, 20% glycerol, 100 mM Tris-HCl pH7.5, 2% Tween 20, 0.5 mM phosphodiesterase inhibitors, and 1 mM PMSF). Proteins were subjected to 5-12% gels and transferred onto PVDF membranes (Cat. #IPFL00010, Millipore, Billerica, MA, USA) at room temperature for 1 hour at 120 V in transfer buffer containing 25 mM Tris, 192 mM glycine, and 20% methanol in milliQ water. The membranes were blocked in 5% nonfat milk blocking buffer for 40 minutes. Samples were incubated with the primary antibodies at 4°C overnight, and then HRP-conjugated secondary antibodies at room temperature for 2 hours. Primary antibodies were used in the following concentrations: mouse anti-α-Tubulin (1:5000, Cat. #T9026, Sigma), rabbit anti-Cathepsin B (1:1000, Cat. #31718S, Cell Signaling), rabbit anti-Cathepsin D (1:1000, Cat. #A19680, ABclonal), rabbit anti-Myc (1:2000, Cat. #0912-2, Hua An Biotechnology), rabbit anti-ATP6V1C1 (1:100, Cat. # A18253, ABclonal), and mouse anti-Flag (1:1000, Cat. #F3165, Sigma, St. Louis, MO, USA). Bands were visualized by Clarity Western ECL Substrate (Cat. #WBKLS0500, Millipore). Results are more than three independent biological replicates. Secondary antibodies were from Jackson ImmunoResearch: goat anti-mouse-HRP (1:5000, Cat. #115-035-003), and goat anti-rabbit-HRP (1:5000, Cat. #111-036-003) and goat anti-rat-HRP (1:5000, Cat. #112-035-003).

Co-immunoprecipitation: S2 cells were transfected and lysed on ice by repeat pipetting in lysis buffer (0.4% NP-40, 0.2 mM EDTA, 150 mM NaCl, 20% glycerol, 100 mM Tris-HCl pH7.5, 2% Tween 20, 0.5 mM phosphodiesterase inhibitors, and 1 mM PMSF) for 30 minutes. Samples were then centrifuged at 13,000 rpm at 4°C for 10 minutes for pull-downs, the soluble supernatant was incubated with prewashed anti-FLAG(R) M2 beads (Cat. #A2220; Sigma) at 4°C overnight. Proteins were subjected to 5-12% gels and transferred onto separated on SDS-PAGE gels and transferred to PVDF membranes (Cat. #IPFL00010, Millipore) at room temperature for 1 hour at 120 V in transfer buffer containing 25 mM Tris, 192 mM glycine, and 20% methanol in milliQ water. Primary antibodies were used in the following concentrations: rabbit anti-ATP6V1C1 (1:100, Cat. #A18253, ABclonal), rat anti-dAux (1:100)^5^, mouse-anti-Flag (1:1000, Cat. #F3165, Sigma) and rabbit-anti-GFP (1:2000, SB-AB0047, ShareBio).

**Behavior analysis**

Mouse footprint gait analysis: To perform footprint gait analysis, mouse hindpaws and forepaws were coated with black and red nontoxic paint, and trained to walk along a 33 cm long and 8 cm wide open-top runway (with 6.35 cm high walls). A fresh sheet of white paper was placed on the floor of the runway for each run. The footprint patterns were assessed quantitatively by stride length^6^.

Mouse pole test: Mice were placed with a 55 cm metal pole of 1 cm diameter covered with adhesive tape to facilitate traction. Upon positioned head-up at the top of the pole, the time required for mice to turn (t-turn) and climb down completely was recorded.

Fly lifespan analysis: To clean the genetic background, *repo-*GAL4, *UAS-LacZ*, and *UAS-daux-*RNAi fly strains were separately back-crossed to isogenic *w^1118^* for five generations. Newly eclosed male or female flies (n=200 for each genotype) were collected and placed separately in 10 vials (each contains 20 unisex flies). Fresh food and new vials were provided every other day. Survival experiments were repeated at least three times. Mortality (live/total, %) was analyzed.

Fly locomotion analysis: The rapid iterative negative geotaxis (RING) assay was used to analyze climbing ability^7-9^. In brief, flies at different age were collected and placed in vials with fly food (no yeast) for 1 day before test (n=100 for each genotype). Cohorts of 100 flies were subjected in ten cylinders (inner diameter: 20 mm; height: 200 mm) placed side by side, and each cylinder contains ten unisex flies. An initial mechanical shock was applied six times, and all flies were tapped down to the bottom of the vertically-positioned tube, then the limbing distances for each fly every 6 seconds were recorded and analyzed by RflyDetection software, which allows automatic record of fly position within the tube using video images (Sony digital camera, HDR-CX220E). Data were from at least three independent experiments.

Fly sleep analysis: The parental and F1 generation flies were raised on standard medium in an incubator set to 25°C with 12-hour light/12-hour dark (LD) cycle. The flies were collected at the age of 0-2 days old and mated for 2 days. Each mated female was then loaded into sleep tubes (PGT5x65 Pyrex Glass, Trikinetics) that contained 2% agar and 5% sucrose at one end. The other end was sealed with parafilm with a hole for air-circulation. The tubes were placed into the DAM2 *Drosophila* activity monitor system (Trikinetics, Waltham) for sleep recording. Activity was recorded for five consecutive days including entrainment for the first two days. Data from the following three days were averaged for further analysis. The original data acquired by the DAM system software were processed by DAMfileScan113. Sleep analysis was then performed using an in-house program called SCAMP2019v2^10^ in Matlab (MathWorks). Parameters of total sleep and sleep latency were calculated.

**Microglia morphology analysis**

The features of microglia morphology were elaborated by Imaris 3D interactive visualization software. Briefly, the immunofluorescent staining for Iba1 was acquired by Nikon TI2-E+CSU W1 Sora Spinning Disk confocal microscope (60X objective NA=1.4, Tokyo, Japan), and each microglial cell was isolated from the context. The resulting image (containing one cell) was submitted to Imaris for the semi-automated detection of processes and terminal points of branches by using the “filament” plug-in. The final 3D rendering surface images were made by combining “surface” and “filament” for the representative image.

**Preparation of α-synuclein preformed fibril (PFF)**

α-synuclein preformed fibril (α-syn PFF) was prepared as previously described^11^. Briefly, pET22 vector carrying full-length wildtype mouse *SNCA* gene was co-expressed with the yeast N-acetyltransferase complex B in BL21 (DE3) cells to obtain the N-terminally acetylated α-syn protein. α-syn proteins were purified using anion exchange column followed by Superdex 75 and then incubated at 37°C in solution (100 μM in 50 mM Tris, pH 7.5, 150 mM KCl, and 0.05% NaN3 buffer) with constant agitation (900 rpm) in ThermoMixer for 7 days. The fibril yield was calculated as the total amount of α-syn monomer subtracting the amount of residual soluble α-syn after pelleting the fibrils. The pelleted fibrils were suspended with PBS to 2 μg/μL. Final α-Syn PFFs were obtained after sonication (20% power, 15 cycles, each cycle contained 1 second on and 1 second off, JY92-IIN sonicator) on ice. The physical state of the fibrils was monitored by TEM.

**Bilateral inoculation of PFF into mouse dorsal striatum**

6 to 8-week-old mice of varied strains were anesthetized with 0.5% isoflurane vapor mixed with 1% O_2_. α-syn PFFs (2 μg/μl) or sterile PBS (0.1 μl/g (body weight)) were stereotaxically injected into dorsal striatum of both hemispheres at a dose of 0.2 μg/g (body weight) using the following coordination: +0.2 mm to bregma, ± 2.0 mm from midline, and −2.6 mm from dura. The coordinates of each mouse were scaled by multiplying the scaling factor ζ (ζ = Measured distance between bregma and lambda(mm) 4.21 mm) to acquire the accurate and consistent injecting location. The injection was done with a 10 μl microsyringe at a speed of 0.5 μl/min for the first 0.2 μl and 0.2 μl/min for the rest. After recovery from the anesthetization, mice were transferred back to their home cages with regular housing conditions until the next experiment.

**Confocal microscopy and statistical analysis**

Serial Z-stack sections images were ­acquired at the similar planes of brains across different genotypes by Nikon C2 or TI2-E+CSU W1 Sora Spinning Disk confocal microscope (20x objective NA=0.75, 60x oil objective NA=1.4, Tokyo, Japan), and representative single layer or maximum projection images were shown. Image J (National Institutes of Health) was used to analyze the number, size, and intensity of puncta and colocalization of different channels. Puncta were auto-selected by adjusting the threshold of the images, and then analyzed with “Analyze Particles” plugin. Colocalization were analyzed automatically by the “Colocalization” plugin, and the results were shown as Pearson’s Correlation (R value, no threshold) or Manders’ Correlation (M1 or M2). DA neurons were manually counted with the “Annotation and Measurement’’ plug-in by Nikon analyzing software.

All statistical graphs were analyzed and displayed by GraphPad Prism 8. For statistical analysis, Shapiro-Wilk normality test was firstly used to check the data distribution, when the data were normally distributed, two-tailed unpaired t-test or ordinary one-way ANOVA followed by Tukey’s multiple comparisons test was used, otherwise, Mann Whitney test or Kruskal-Wallis tests followed by Dunn’s multiple comparisons test were used. P value less than 0.05 is considered significant. ns: no significance, p≥ 0.05; *: p<0.05; **: p<0.01; ***: p<0.001; ****: p<0.0001.

**References**

1.	Yang M, Wang H, Chen C, Zhang S, Wang M, Senapati B, et al. Glia-derived temporal signals orchestrate neurogenesis in the Drosophila mushroom body. Proc Natl Acad Sci U S A 2021; 118.

2.	Pulipparacharuvil S, Akbar MA, Ray S, Sevrioukov EA, Haberman AS, Rohrer J, Kramer H. Drosophila Vps16A is required for trafficking to lysosomes and biogenesis of pigment granules. J Cell Sci 2005; 118:3663-73.

3.	McCarthy RC, Lu DY, Alkhateeb A, Gardeck AM, Lee CH, Wessling-Resnick M. Characterization of a novel adult murine immortalized microglial cell line and its activation by amyloid-beta. J Neuroinflammation 2016; 13:21.

4.	Li H, Li Y, Lei Z, Wang K, Guo A. Transformation of odor selectivity from projection neurons to single mushroom body neurons mapped with dual-color calcium imaging. Proc Natl Acad Sci U S A 2013; 110:12084-9.

5.	Zhang S, Yi S, Wang L, Li S, Wang H, Song L, et al. Cyclin-G-associated kinase GAK/dAux regulates autophagy initiation via ULK1/Atg1 in glia. Proc Natl Acad Sci U S A 2023; 120:e2301002120.

6.	Wertman V, Gromova A, La Spada AR, Cortes CJ. Low-Cost Gait Analysis for Behavioral Phenotyping of Mouse Models of Neuromuscular Disease. J Vis Exp 2019.

7.	Gargano JW, Martin I, Bhandari P, Grotewiel MS. Rapid iterative negative geotaxis (RING): a new method for assessing age-related locomotor decline in Drosophila. Exp Gerontol 2005; 40:386-95.

8.	Cao W, Song L, Cheng J, Yi N, Cai L, Huang FD, Ho M. An Automated Rapid Iterative Negative Geotaxis Assay for Analyzing Adult Climbing Behavior in a Drosophila Model of Neurodegeneration. J Vis Exp 2017.

9.	Song L, He Y, Ou J, Zhao Y, Li R, Cheng J, et al. Auxilin Underlies Progressive Locomotor Deficits and Dopaminergic Neuron Loss in a Drosophila Model of Parkinson's Disease. Cell reports 2017; 18:1132-43.

10.	Donelson NC, Kim EZ, Slawson JB, Vecsey CG, Huber R, Griffith LC. High-resolution positional tracking for long-term analysis of Drosophila sleep and locomotion using the "tracker" program. PLoS One 2012; 7:e37250.

11.	Long H, Zheng W, Liu Y, Sun Y, Zhao K, Liu Z, et al. Wild-type alpha-synuclein inherits the structure and exacerbated neuropathology of E46K mutant fibril strain by cross-seeding. Proc Natl Acad Sci U S A 2021; 118.

**Supplementary Figures**

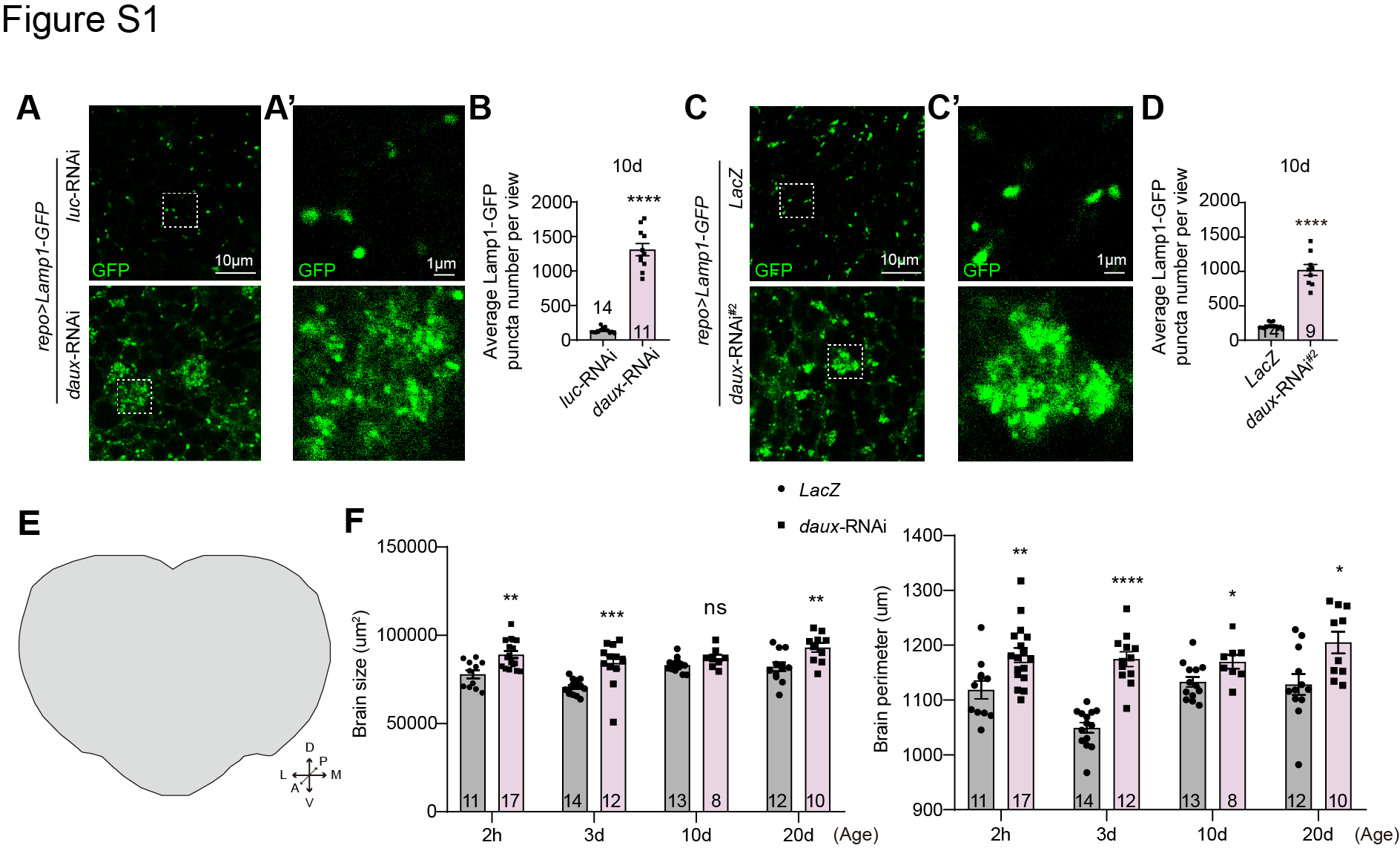

**Figure S1 (related to Figure 1). Lack of dAux increases the glial lysosome number and brain size** (**A**-**D**) The number of lysosomes in 10-day-old adult fly brains were analyzed by comparing to a different control *UAS-luciferase*-RNAi (*luc*-RNAi, A and B) or a second RNAi line, *UAS-daux*-RNAi^#2^ (BL#39017, C and D). Note that results are consistent with the main Figures when using *UAS-LacZ* as a control or the other *daux*-RNAi throughout the study. (**E**) An illustration of an adult fly brain for size and perimeter analysis. (**F**) The overall brain size and perimeter of flies lacking glial dAux are larger and longer, respectively at different ages. Scale bars of different sizes are indicated on the images. Serial confocal Z-stack sections were taken at similar planes across all genotypes, with representative images shown as single layers. Quantification Figures from the same images but with different parameters are shown under the same label unless noted otherwise. Statistical graphs are shown with scatter dots indicating the number of brain samples analyzed (n, also on the bar). Data are shown as mean ± SEM. P-values of significance (indicated with asterisks, ns no significance, * p<0.05, ** p<0.01, *** p<0.001, and **** p<0.0001) are calculated by two-tailed unpaired t-test or Mann Whitney test.

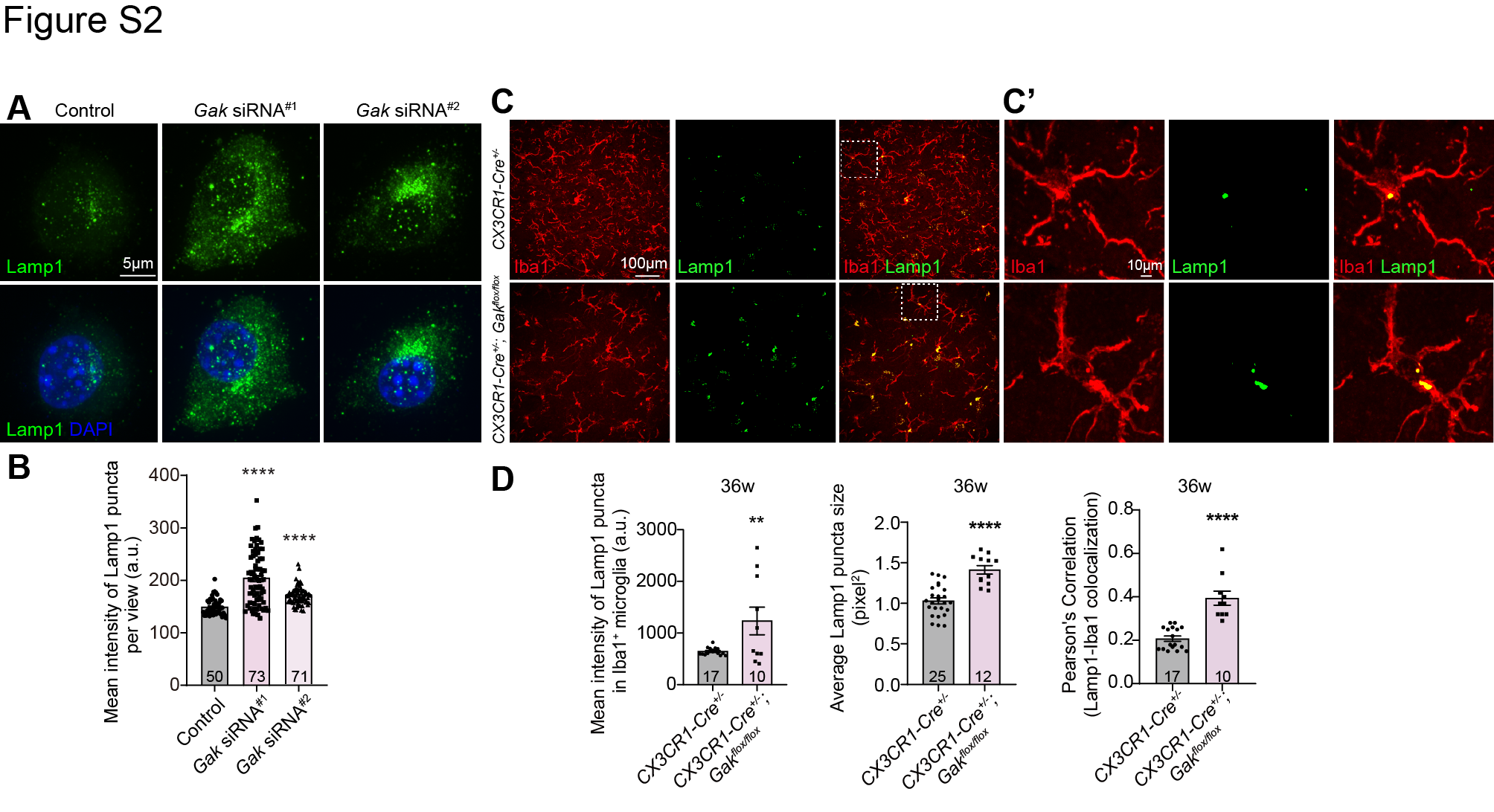

**Figure S2 (related to Figure 1). Lack of Gak increases the lysosome number in immortalized microglia and mouse primary microglia** (**A** and **B**) Representative images (A) and quantifications (B) of lysosomes in IMG cells. Note that the intensities of the Lamp1-positive lysosomes (green) increase upon Gak depletion by *Gak* siRNAs. (**C** and **D**) Representative images (C) and quantifications (D) of glial lysosomes in mouse primary microglia in substantia nigra. The areas in white dotted squares in C are enlarged, aligning on the right (C’). Note that the intensities and size of Lamp1-positive lysosomes (green) in Iba1-positive microglia and the Lamp1-Iba1 colocalization increase in the *Gak* cKO mice at 36 weeks (D). Scale bars of different sizes are indicated on the images. Serial confocal Z-stack sections were taken at the similar planes across all genotypes, with representative images shown as maximal projection. Quantification Figures from the same images but with different parameters are shown under the same label unless noted otherwise. Colocalization is analyzed using the Pearson’s Correlation. Statistical graphs are shown with scatter dots indicating the cell numbers analyzed (n, also on the bar). Data are shown as mean ± SEM. P-values of significance (indicated with asterisks, ns no significance, * p<0.05, ** p<0.01, *** p<0.001, and **** p<0.0001) are calculated by two-tailed unpaired t-test, Mann Whitney test, or Kruskal-Wallis tests followed by Dunn’s multiple comparisons test.

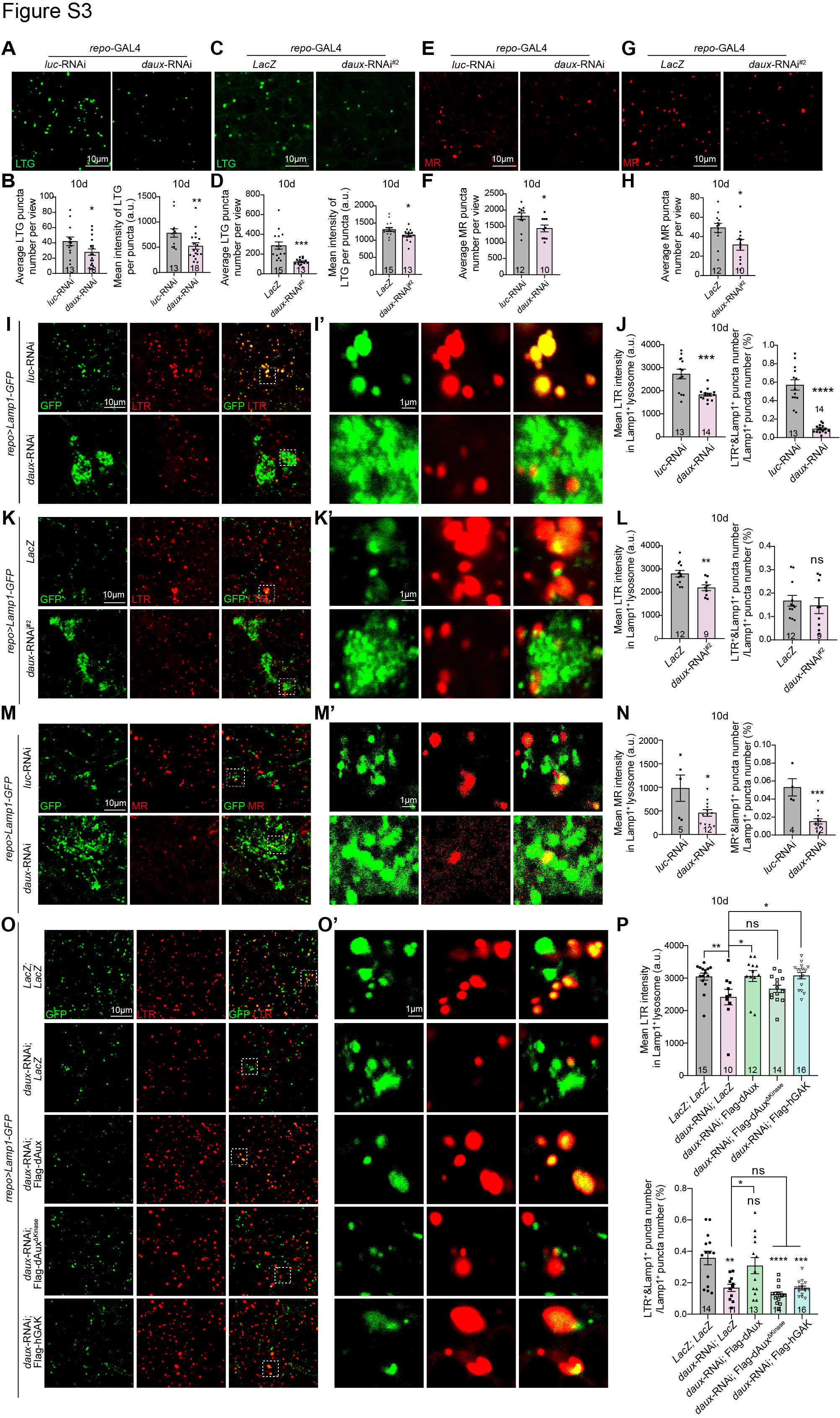

**Figure S3 (related to Figure 1). Lack of dAux disrupts lysosomal acidification in glia (A**-**H)** Representative images (A,C,E,G) and quantifications (B,D,F,H) of LTG- or MR-positive puncta in 10-day-old adult fly brains expressing *daux*-RNAi in glia. The intensities of LTG- (A-D) or MR-positive (E-H) puncta in the whole brain decrease upon *daux*-RNAi expression in glia. **(I**-**N)** Representative images (I,I’,K,K’,M,M’) and quantifications (J,L,N) of LTR- (I-L) or MR-positive (M and N) puncta colocalizing to the glial Lamp1-positive lysosomes in 10-day-old adult fly brains expressing *daux*-RNAi in glia. The areas in white dotted squares in I,K,M are enlarged, aligning on the right (I’,K’,M’). The mean intensities of LTR- or MR- per glial Lamp1-positive lysosomes decrease upon *daux*-RNAi expression in glia. Note that results are consistent to the main Figures when using *UAS-LacZ* as a control or the other *daux*-RNAi throughout the study. (**O** and **P**) Representative images (O,O’) and quantifications (P) of glial lysosomes (Lamp1-GFP, green) staining with LTR dye in 10-day-old adult fly brains with the indicated genotypes. Note that expressing Flag-dAux or Flag-hGAK fully rescues the *daux*-RNAi-induced decrease of LTR intensities in glial Lamp1-positive lysosomes, whereas expressing Flag-dAux^ΔKinase^ fails to do so. The areas in white dotted squares in O are enlarged, aligning on the right (O’). Scale bars of different sizes are indicated on the images. Serial confocal Z-stack sections were taken at the similar planes across all genotypes, with representative images shown as single layer. Quantification Figures from the same images but with different parameters are shown under the same label unless noted otherwise. Statistical graphs are shown with scatter dots indicating the number of brain samples analyzed (n, also on the bar). Data are shown as mean ± SEM. P-values of significance (indicated with asterisks, ns no significance, * p<0.05, ** p<0.01, *** p<0.001, and **** p<0.0001) are calculated by two-tailed unpaired t-test, Mann Whitney test, ordinary one-way ANOVA followed by Tukey’s multiple comparisons test, or Kruskal-Wallis tests followed by Dunn’s multiple comparisons test.

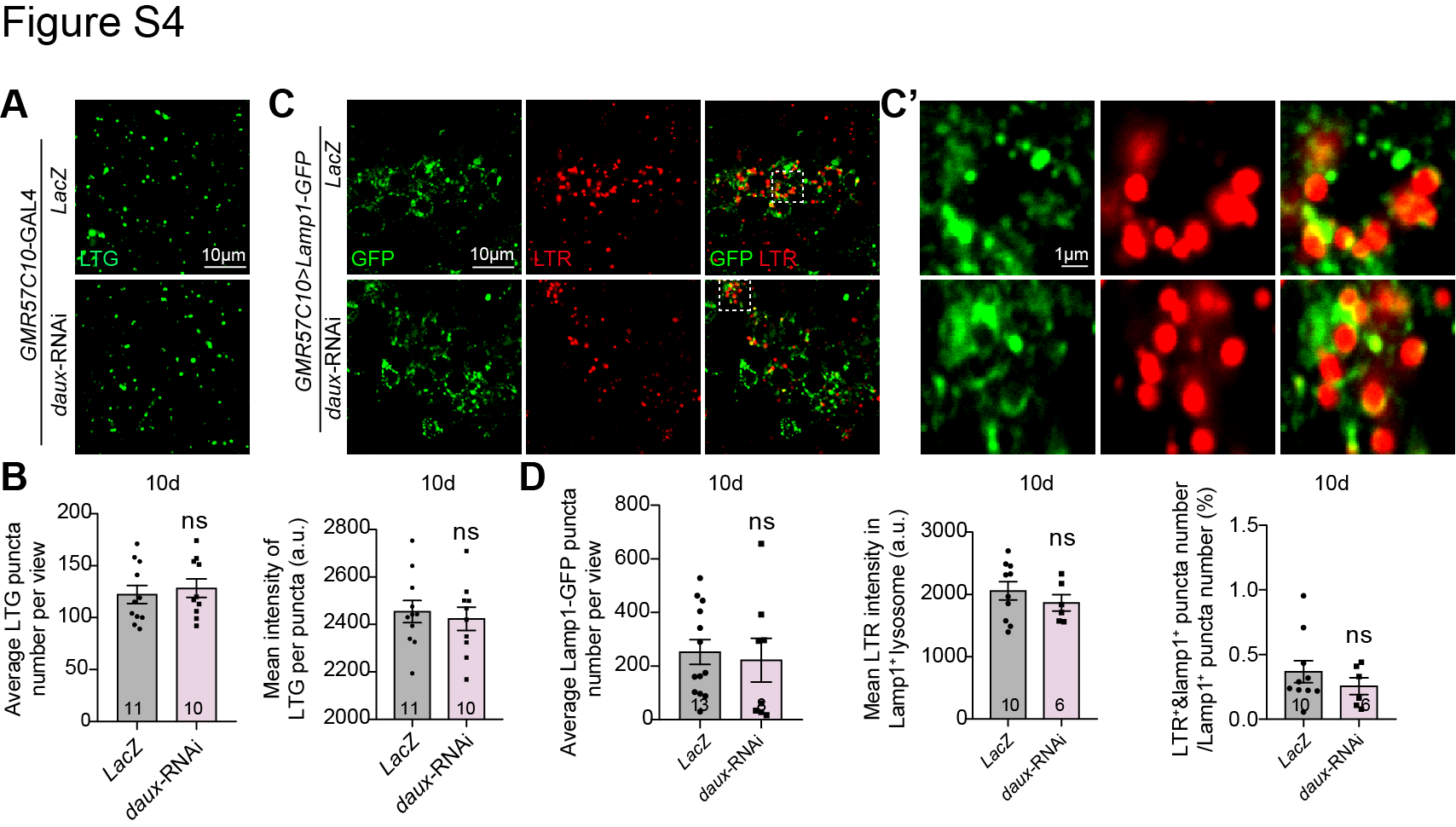

**Figure S4 (related to Figure 1). Lack of dAux does not affect lysosomal acidification in neurons** (**A**-**D**) Representative images (A,C) and quantifications (B,D) of LTG- or LTR-positive puncta in 10-day-old adult fly brains expressing *daux*-RNAi in neurons under the control of the pan-neuronal driver *GMR57C10*-GAL4. Note that both the number and intensities of LTG- (A,B) or LTR-positive (C,D) puncta remain unaffected in the whole brain or per neuronal Lamp1-positive lysosome (green) upon neuronal dAux depletion. The areas in white dotted squares in C are enlarged, aligning on the right (C’). Scale bars of different sizes are indicated on the images. Serial confocal Z-stack sections were taken at the similar planes across all genotypes, with representative images shown as single layer. Quantification Figures from the same images but with different parameters are shown under the same label unless noted otherwise. Statistical graphs are shown with scatter dots indicating the number of brain samples analyzed (n, also on the bar). Data are shown as mean ± SEM. P-values of significance (indicated with asterisks, ns no significance, * p<0.05, ** p<0.01, *** p<0.001, and **** p<0.0001) are calculated by two-tailed unpaired t-test or Mann Whitney test.

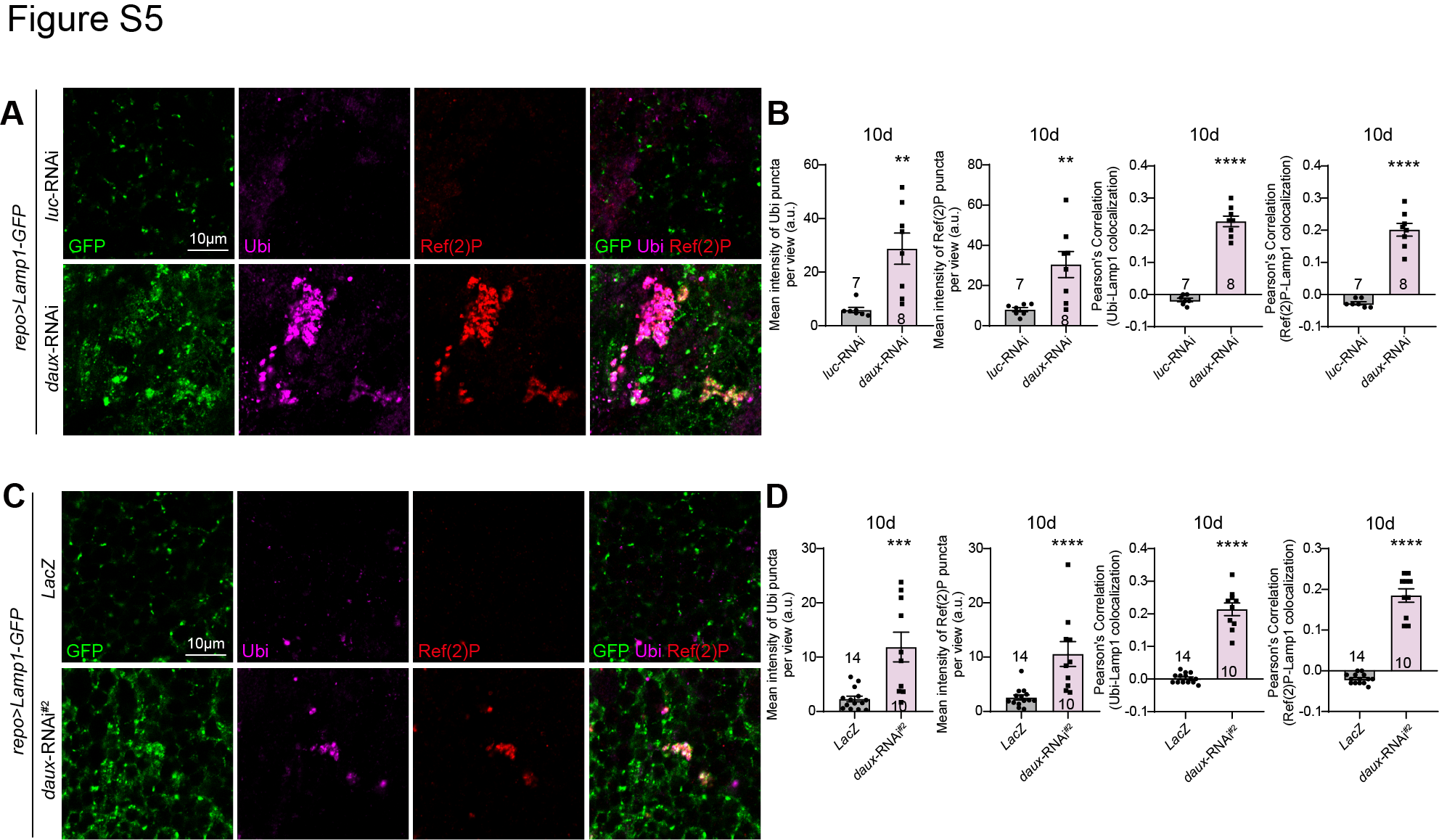

**Figure S5 (related to Figure 2). Lack of dAux causes accumulation of Ref(2)P and Ubi** (**A**-**D**) Representative images (A,C) and quantifications (B,D) of Ubiquitin (Ubi) and Ref(2)P levels in 10-day-old adult fly brains expressing *daux-*RNAi in glia. Note that both the intensities of Ubi and Ref(2)P and the colocalization of either Ubi or Ref(2)P with Lamp1 increase upon glial dAux depletion. Results are consistent to the main Figures when using *UAS-LacZ* as a control or the other *daux*-RNAi throughout the study. Scale bars of different sizes are indicated on the images. Serial confocal Z-stack sections were taken at the similar planes across all genotypes, with representative images shown as single layer. Quantification Figures from the same images but with different parameters are shown under the same label unless noted otherwise. Colocalization is analyzed using the Pearson’s Correlation. Statistical graphs are shown with scatter dots indicating the number of brain samples analyzed (n, also on the bar). Data are shown as mean ± SEM. P-values of significance (indicated with asterisks, ns no significance, * p<0.05, ** p<0.01, *** p<0.001, and **** p<0.0001) are calculated by two-tailed unpaired t-test or Mann Whitney test.

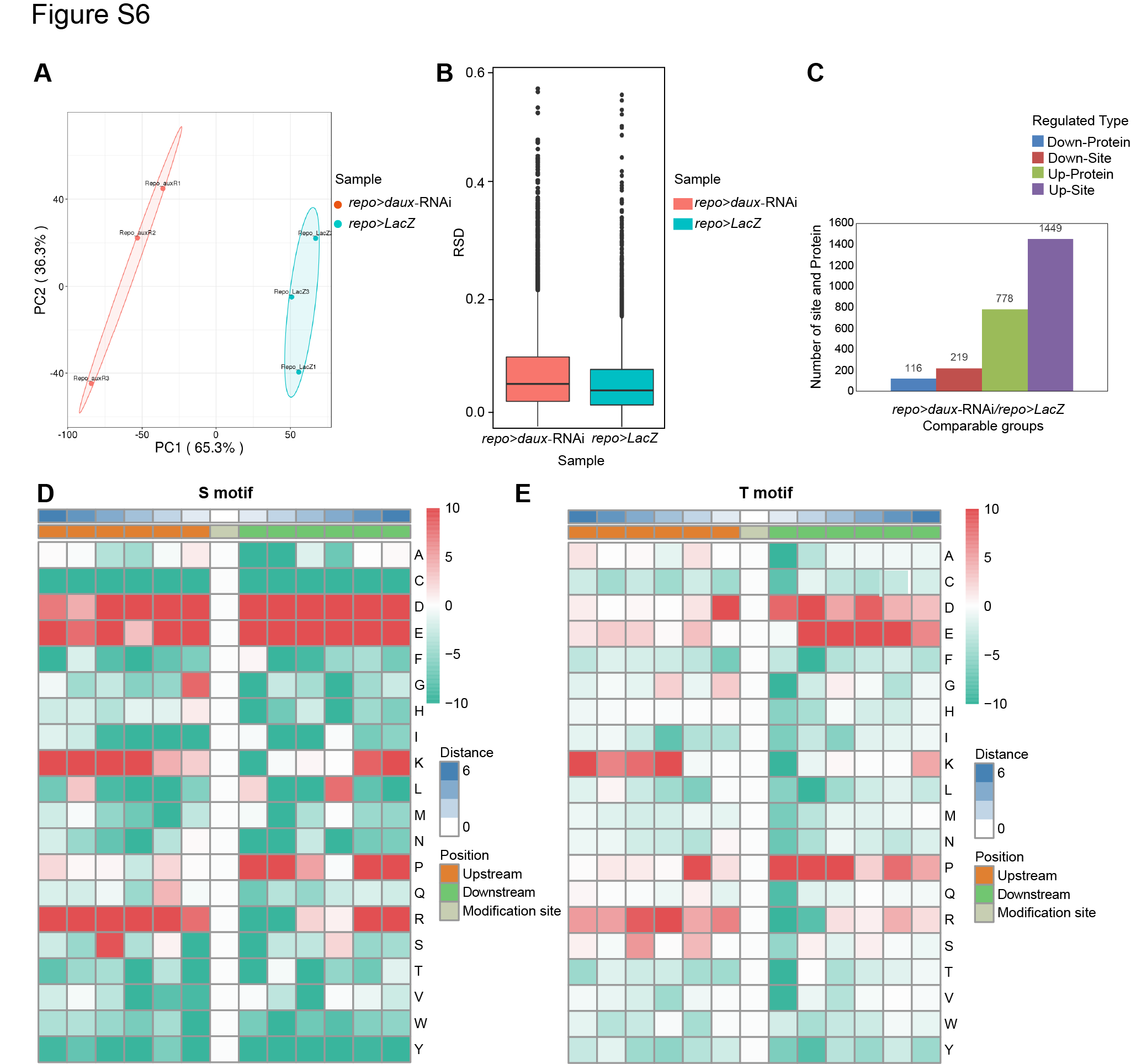

**Figure S6 (related to Figure 3). Phosphoproteomic analysis identifying potential dAux substrates** (**A** and **B**) Principal component analysis (PCA) and relative standard deviation (RSD) are used to assess protein quantification reproducibility (n=3 per genotype). Note that the quantitative results of biological replicates or technical replicates are statistically consistent. (**C**) The total number of modified proteins and sites collected from adult fly brains expressing *daux*-RNAi in glia. A change larger than 1.3 is defined as upregulation, and a change less than 0.77 (1/1.3) is defined as downregulation, respectively. (**D** and **E**) Heatmap of motif enrichment upstream and downstream of S (D) or T (E) phosphorylation sites. Red represents significant enrichment near the phosphorylation site, and green represents significant decrease near the phosphorylation site.

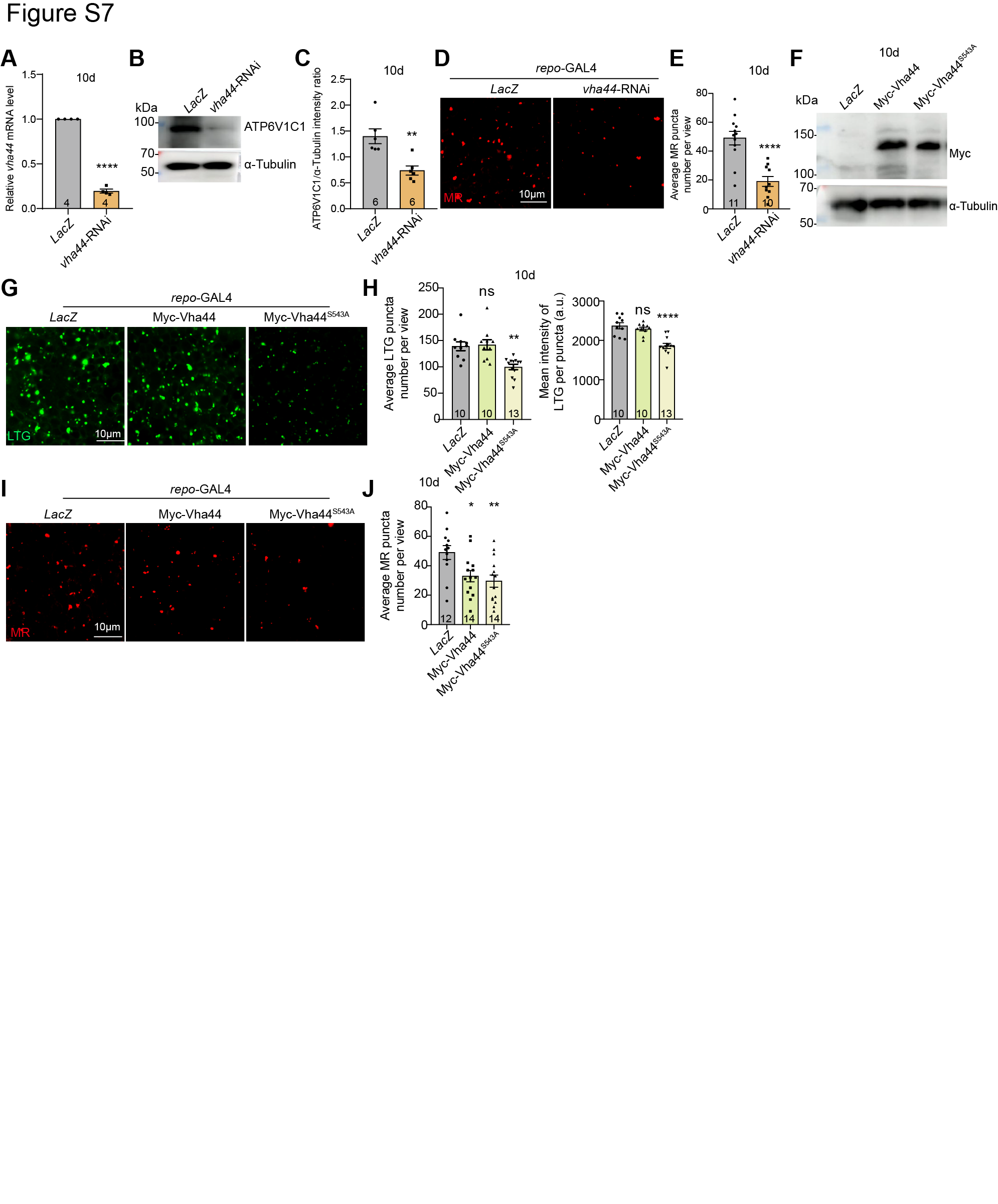

**Figure S7 (related to Figure 4) dAux-mediated Vha44 S543 phosphorylation regulates lysosomal acidification in glia** (**A-C**) The knockdown efficiency of *vha44*-RNAi was analyzed by qRT-PCR (A) and WB (C and D) analyses. RNA and protein samples were collected from the 10-day-old adult fly heads (control and *repo>vha44-*RNAi). Note that both *vha44* mRNA and protein levels decrease when expressing *vha44*-RNAi in glia. *rp49* or α-Tubulin serve as controls for qRT-PCR and WB, respectively. (**D** and **E**) Representative images (D) and quantifications (E) of MR staining in the 10-day-old adult fly brains. Note that downregulating glial *vha44* reduces the number of MR-positive puncta. **(F)** Expressions of wild-type Myc-Vha44 and Myc-Vha44^S543A^ mutant are validated by WB using samples collected from the 10-day-old adult fly heads. (**G** and **H**) Representative images (G) and quantification (H) of LTG-positive puncta in 10-day-old adult fly brains expressing Myc-Vha44 and Myc-Vha44^S543A^. Note that the number and intensities of LTG-positive puncta decrease upon Myc-Vha44^S543A^ expression. **(I** and **J)** Representative images (I) and quantifications (J) of MR staining in 10-day-old adult fly brains. Note that the number of MR-positive puncta decreases more significantly when expressing Myc-Vha44^S543A^ in glia. Scale bars are indicated on the images. Serial confocal Z-stack sections were taken at the similar planes across all genotypes, with representative images shown as single layer. WB statistics are done with at least three independent biological replicates. Quantification Figures from the same images but with different parameters are shown under the same label unless noted otherwise. Statistical graphs are shown with scatter dots indicating the number of brain samples or biological replicates analyzed (n, also on the bar). Data are shown as mean ± SEM. P-values of significance (indicated with asterisks, ns no significance, * p<0.05, ** p<0.01, *** p<0.001, and **** p<0.0001) are calculated by two-tailed unpaired t-test, Mann Whitney test, ordinary one-way ANOVA followed by Tukey’s multiple comparisons test, or Kruskal-Wallis tests followed by Dunn’s multiple comparisons test.

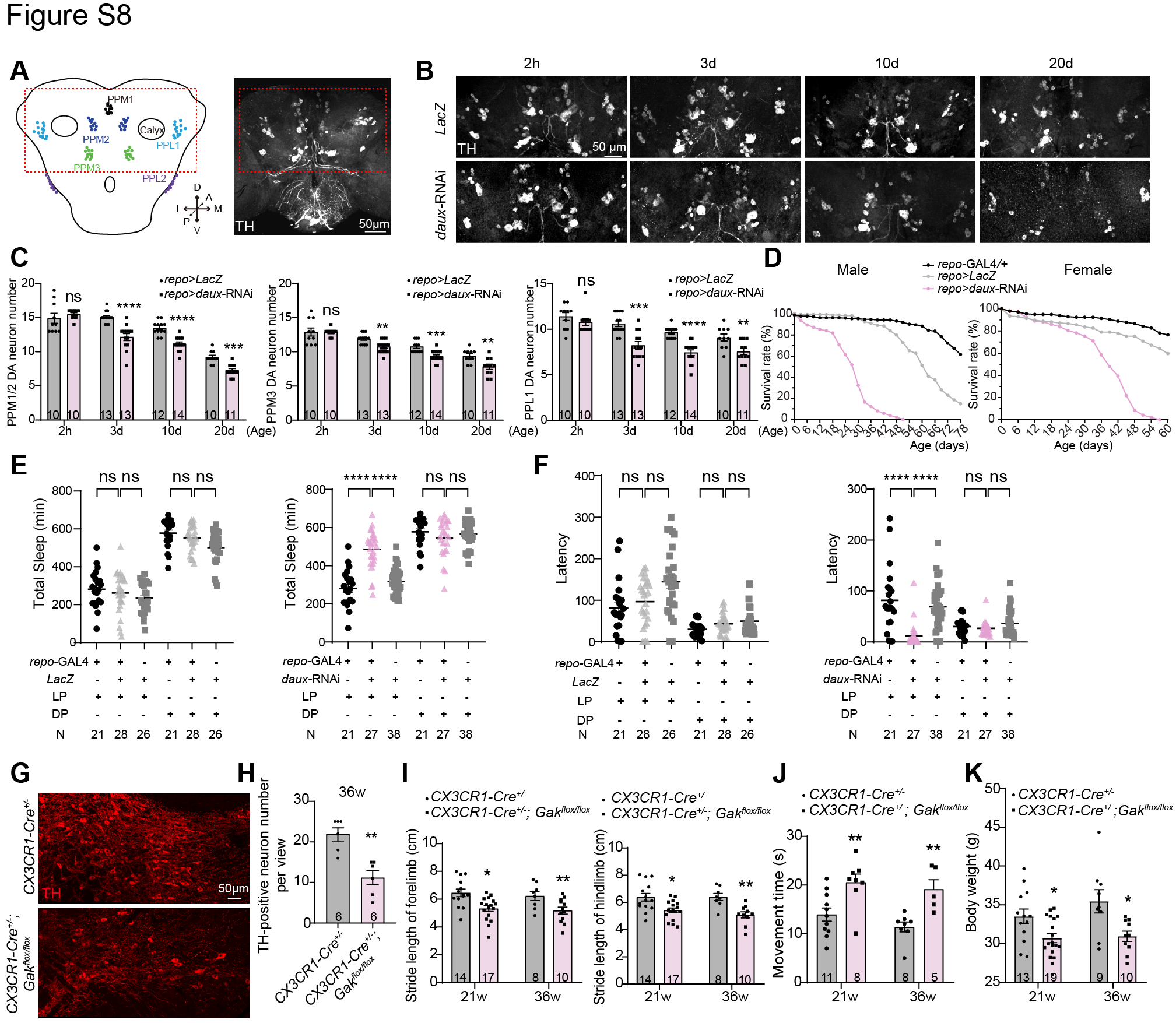

**Figure S8 (related to Figure 6 and Figure 7). Lack of glial Gak/dAux leads to a broad spectrum of symptoms implicated in PD** (**A**) An illustration of the DA neuron clusters in the adult fly brains. These clusters include protocerebral posterior lateral (PPL)1, PPL2, protocerebral posterior medial (PPM)1, PPM2, and PPM3. An acquired microscopic image of an adult fly brain stained with anti-TH antibodies revealing all DA neuron clusters. The adult fly brain is positioned with the coordinates described (A: Anterior, P: Posterior, M: Medial, L: Lateral, D: Dorsal, V: Ventral), with the region enclosed by red dotted squares selected for imaging. (**B** and **C**) Representative images (B) and quantifications (C) of DA neuron number in the control and *repo*>*daux*-RNAi adult fly brains. Note that DA neuron number in the PPM1/2, PPM3, and PPL1 clusters decreases in an age-dependent manner upon glial dAux depletion. At 2 hours after eclosion, the DA neuron number remains unaffected, ruling out the developmental defects. (**D**) The lifespan of both male and female flies is shortened upon glial dAux depletion. (**E** and **F**) Lack of glial dAux causes an increase of the total sleep time and a decrease of the sleep latency. (**G** and **H**) The number of DA neurons marked by the anti-TH antibodies in the substantia nigra decreases in microglial *Gak* cKO mice (*CX3CR1-Cre^+/-^; Gak^flox/flox^*) at 36 weeks. (**I**-**K**) Microglial *Gak* cKO mice exhibit locomotor deficits and weight loss at 21 and 36 weeks by footprint analysis, pole test and examination of the body weight. Scale bars are indicated on the images. Serial confocal Z-stack sections were acquired at the similar planes across all genotypes, with representative images shown as maximal projection. Statistical graphs are shown with scatter dots indicating the number of brain samples or fly numbers analyzed (n, also on the bar). In lifespan experiments, n=200 for each genotype. Data are shown as mean ± SEM. P-values of significance (indicated with asterisks, ns no significance, * p<0.05, ** p<0.01, *** p<0.001, and **** p<0.0001) are calculated by two-tailed unpaired t-test, Mann Whitney test, ordinary one-way ANOVA followed by Tukey’s multiple comparisons test, or Kruskal-Wallis tests followed by Dunn’s multiple comparisons test.

**Table S1**

Detailed fly or mouse genotypes in each experiment categorized by Figures.

| Figure | Genotype |
| --- | --- |
| Fig. 1 | |
| (B) | *UAS-Lamp1-GFP/+; repo-*GAL4*/+* |
| (C and D) | *UAS-Lamp1-GFP/+; repo-*GAL4*/UAS-LacZ* |
|  | *UAS-Lamp1-GFP/UAS-daux-*RNAi*; repo-*GAL4*/+* |
| (E) | *UAS-HRP/+; repo-*GAL4*/+* |
| (F and G) | *UAS-HRP/+; repo-*GAL4*/UAS-LacZ* |
|  | *UAS-HRP/UAS-daux-*RNAi*; repo-*GAL4*/+* |
| (H-O) | *repo-*GAL4*/UAS-LacZ* |
|  | *UAS-daux-*RNAi*/+; repo-*GAL4*/+* |
| (P-S) | *UAS-Lamp1-GFP/+; repo-*GAL4*/UAS-LacZ* |
|  | *UAS-Lamp1-GFP/UAS-daux-*RNAi*; repo-*GAL4*/+* |
| Fig. 2 | |
| (A and B) | *UAS-Lamp1-GFP/+; repo-*GAL4*/UAS-LacZ* |
|  | *UAS-Lamp1-GFP/UAS-daux-*RNAi*; repo-*GAL4*/+* |
| (C and D) | *UAS-LacZ/+; repo-*GAL4*, UAS-mCD8-RFP, GMR57C10-*lexA*/lexAop2-6xMyc-SNCA* |
|  | *UAS-daux-*RNAi*/+; repo-*GAL4*, UAS-mCD8-RFP, GMR57C10-*lexA*/lexAop2-6xMyc-SNCA* |
| (E and F) | *UAS-LacZ/UAS-mCherry-Lamp1; repo-*GAL4*, GMR57C10-*lexA*/lexAop2-6xMyc-SNCA* |
|  | *UAS-daux-*RNAi*/UAS-mCherry-Lamp1; repo-*GAL4*, GMR57C10-*lexA*/lexAop2-6xMyc-SNCA* |
| Fig. 3 | |
| (B-E) | *repo-*GAL4*/UAS-LacZ* |
|  | *UAS-daux-*RNAi*/+; repo-*GAL4*/+* |
| (I and J) | *repo-*GAL4*/UAS-LacZ* |
|  | *UAS-daux-*RNAi*/+; repo-*GAL4*/+* |
| Fig. 4 | |
| (A and B) | *repo-*GAL4*/UAS-LacZ* |
|  | *repo-*GAL4*/UAS-vha44-*RNAi |
| (C and D) | *UAS-Lamp1-GFP/+; repo-*GAL4*/UAS-LacZ* |
|  | *UAS-Lamp1-GFP/+; repo-*GAL4*/UAS-vha44-*RNAi |
| (E-H) | *UAS-LacZ/+; repo*-GAL4*/UAS-LacZ* |
|  | *UAS-daux-*RNAi*/+; repo-*GAL4*/UAS-LacZ* |
|  | *UAS-daux-*RNAi*/+; repo*-GAL4*/UAS-6xMyc-Vha44* |
|  | *UAS-daux-*RNAi*/+; repo-*GAL4*/UAS-6xMyc-Vha44^S543A^* |
| (I-L) | *UAS-Lamp1-GFP/UAS-LacZ; repo-GAL4/UAS-LacZ* |
|  | *UAS-Lamp1-GFP/UAS-daux-*RNAi*; repo-*GAL4*/UAS-LacZ* |
|  | *UAS-Lamp1-GFP/UAS-daux-*RNAi*; repo-*GAL4*/UAS-6xMyc-Vha44* |
|  | *UAS-Lamp1-GFP/UAS-daux-*RNAi*; repo-*GAL4*/UAS-6xMyc-Vha44^S543A^* |
| Fig. 5 | |
| (B, C, F-I) | *UAS-mCherry-Vha100-2/UAS-LacZ; repo-*GAL4*/UAS-EGFP-Vha44* |
|  | *UAS-mCherry-Vha100-2/UAS-daux-*RNAi*; repo-*GAL4*/UAS-EGFP-Vha44* |
|  | *UAS-mCherry-Vha100-2/UAS-LacZ; repo-*GAL4*/UAS-EGFP-Vha44^S543A^* |
|  | *UAS-mCherry-Vha100-2/UAS-daux-*RNAi*; repo-*GAL4*/UAS-EGFP- Vha44^S543A^* |
| (D and E) | *UAS-mCherry-Lamp1/UAS-LacZ; repo-*GAL4*/UAS-EGFP-Vha44* |
|  | *UAS-mCherry-Lamp1/UAS-daux-*RNAi*; repo-*GAL4*/UAS-EGFP-Vha44* |
|  | *UAS-mCherry-Lamp1/UAS-LacZ; repo-*GAL4*/UAS-EGFP-Vha44^S543A^* |
|  | *UAS-mCherry-Lamp1/UAS-daux-*RNAi*; repo-*GAL4*/UAS-EGFP- Vha44^S543A^* |
| Fig. 6 | |
| (B, C, and H) | *repo-*GAL4*/UAS-LacZ* |
|  | *repo-*GAL4*/UAS-vha44-*RNAi |
| (D, E and I) | *repo-*GAL4*/UAS-LacZ* |
|  | *repo-*GAL4*/UAS-6xMyc-Vha44* |
|  | *repo-*GAL4*/UAS-6xMyc-Vha44^S543A^* |
| (F, G, and J) | *UAS-LacZ/+; repo-*GAL4*/UAS-LacZ* |
|  | *UAS-daux-*RNAi*/+; repo-*GAL4*/UAS-LacZ* |
|  | *UAS-daux-*RNAi*/+; repo-*GAL4*/UAS-6xMyc-Vha44* |
|  | *UAS-daux-*RNAi*/+; repo*-GAL4*/UAS-6xMyc-Vha44^S543A^* |
| Fig. 7 | |
| (A-K) | *CX3CR1-Cre^+/-^* |
|  | *CX3CR1-Cre^+/-^; Gak^flox/flox^* |
| Fig. S1 | |
| (A and B) | *UAS-Lamp1-GFP/+; repo-*GAL4*/UAS-luc-*RNAi |
|  | *UAS-Lamp1-GFP/UAS-daux*-RNAi*; repo-*GAL4*/+* |
| (C and D) | *UAS-Lamp1-GFP/+; repo-*GAL4*/UAS-LacZ* |
|  | *UAS-Lamp1-GFP/UAS-daux-*RNAi^#2^*; repo-*GAL4*/+* |
| (F) | *repo-*GAL4*/UAS-LacZ* |
|  | *UAS-daux-*RNAi*/+; repo-*GAL4*/+* |
| Fig. S2 | |
| (C and D) | *CX3CR1-Cre^+/-^* |
|  | *CX3CR1-Cre^+/-^; Gak^flox/flox^* |
| Fig. S3 | |
| (A, B, E and F) | *repo-*GAL4*/UAS-luc-*RNAi |
|  | *UAS-daux-*RNAi*/+; repo-*GAL4*/+* |
| (C, D, G and H) | *repo-*GAL4*/UAS-LacZ* |
|  | *UAS-daux-*RNAi^#2^*/+; repo-*GAL4*/+* |
| (I, J, M and N) | *UAS-Lamp1-GFP/+; repo-*GAL4*/UAS-luc-*RNAi |
|  | *UAS-Lamp1-GFP/UAS-daux*-RNAi*; repo-*GAL4*/+* |
| (K and L) | *UAS-Lamp1-GFP/+; repo-*GAL4*/UAS-LacZ* |
|  | *UAS-Lamp1-GFP/UAS-daux-*RNAi^#2^*; repo-*GAL4*/+* |
| (O and P) | *UAS-Lamp1-GFP/UAS-LacZ; repo-*GAL4*/UAS-LacZ* |
|  | *UAS-Lamp1-GFP/UAS-daux-*RNAi*; repo-*GAL4*/UAS-LacZ* |
|  | *UAS-Lamp1-GFP/UAS-daux-*RNAi*; repo-*GAL4*/UAS-3xFlag-dAux* |
|  | *UAS-Lamp1-GFP/UAS-daux-*RNAi*; repo-*GAL4*/UAS-3xFlag-dAux^ΔKinase^* |
|  | *UAS-Lamp1-GFP/UAS-daux*-RNAi*; repo-*GAL4*/UAS-3xFlag-hGAK* |
| Fig. S4 | |
| (A and B) | *GMR57C10-*GAL4*/UAS-LacZ* |
|  | *UAS-daux-*RNAi*/+; GMR57C10-*GAL4*/+* |
| (C and D) | *UAS-Lamp1-GFP/+; GMR57C10-*GAL4*/UAS-LacZ* |
|  | *UAS-Lamp1-GFP/UAS-daux-*RNAi*; GMR57C10-*GAL4*/+* |
| Fig. S5 | |
| (A and B) | *UAS-Lamp1-GFP/+; repo-*GAL4*/UAS-luc-*RNAi |
|  | *UAS-Lamp1-GFP/UAS-daux-*RNAi*; repo-*GAL4*/+* |
| (C and D) | *UAS-Lamp1-GFP/+; repo-*GAL4*/UAS-LacZ* |
|  | *UAS-Lamp1-GFP/UAS-daux-*RNAi^#2^*; repo-*GAL4*/+* |
| Fig. S6 | |
| (A-E) | *repo-*GAL4*/UAS-LacZ* |
|  | *UAS-daux-*RNAi*/+; repo-*GAL4*/+* |
| Fig. S7 | |
| (A-E) | *repo-*GAL4*/UAS-LacZ* |
|  | *repo-*GAL4*/UAS-vha44-*RNAi |
| (F-J) | *repo-*GAL4*/UAS-LacZ* |
|  | *repo-*GAL4*/UAS-6xMyc-Vha44* |
|  | *repo-*GAL4*/UAS-6xMyc-Vha44^S543A^* |
| Fig. S8 | |
| (B-C) | *repo-*GAL4*/UAS-LacZ* |
|  | *UAS-daux-*RNAi*/+; repo-*GAL4*/+* |
| (D) | *repo-*GAL4*/+* |
|  | *repo-*GAL4*/UAS-LacZ* |
|  | *UAS-daux-*RNAi*/+; repo-*GAL4*/+* |
| (E and F) | *repo-*GAL4*/UAS-LacZ* |
|  | *UAS-daux-*RNAi*/+; repo-*GAL4*/+* |
| (G-K) | *CX3CR1-Cre^+/-^* |
|  | *CX3CR1-Cre^+/-^; Gak^flox/flox^* |
