## Supplementary Video for "dAux orchestrates the phosphorylation-dependent assembly of the lysosomal V-ATPase in glia and contributes to α-synuclein degradation"

### Slide 1
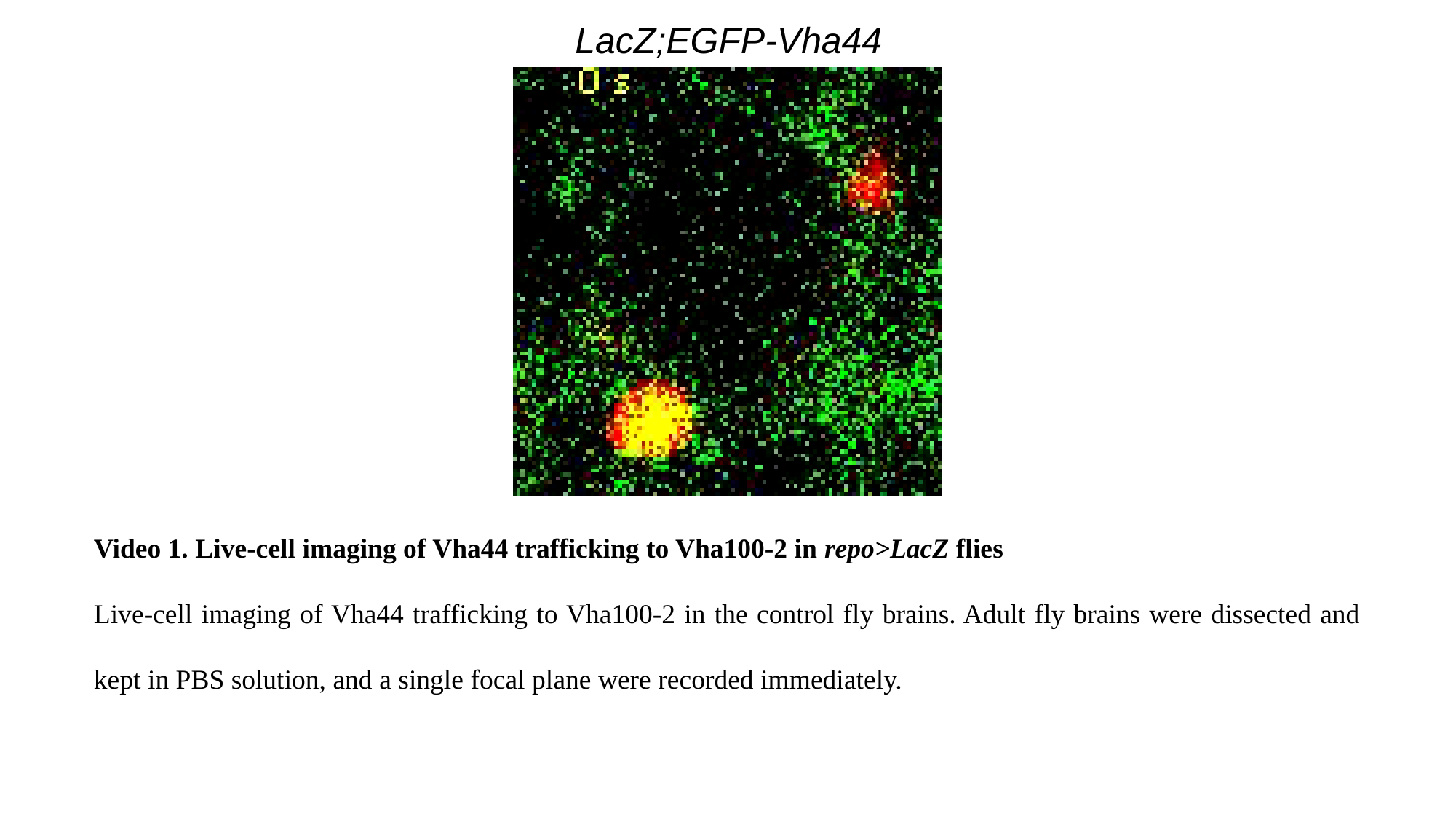

LacZ;EGFP-Vha44
Video 1. Live-cell imaging of Vha44 trafficking to Vha100-2 in repo>LacZ flies
Live-cell imaging of Vha44 trafficking to Vha100-2 in the control fly brains. Adult fly brains were dissected and kept in PBS solution, and a single focal plane were recorded immediately.

### Slide 2
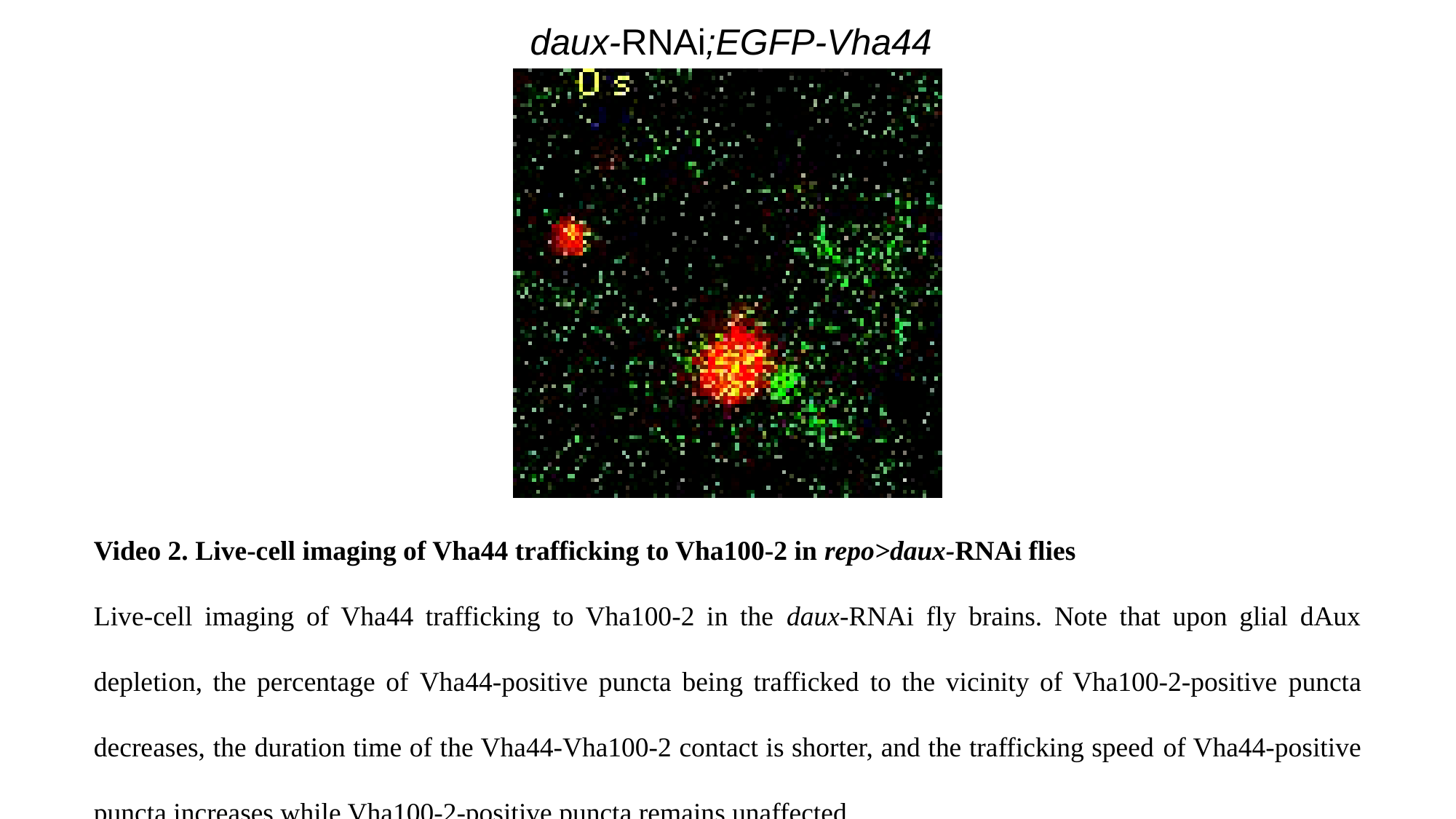

daux-RNAi;EGFP-Vha44
Video 2. Live-cell imaging of Vha44 trafficking to Vha100-2 in repo>daux-RNAi flies
Live-cell imaging of Vha44 trafficking to Vha100-2 in the daux-RNAi fly brains. Note that upon glial dAux depletion, the percentage of Vha44-positive puncta being trafficked to the vicinity of Vha100-2-positive puncta decreases, the duration time of the Vha44-Vha100-2 contact is shorter, and the trafficking speed of Vha44-positive puncta increases while Vha100-2-positive puncta remains unaffected.

### Slide 3
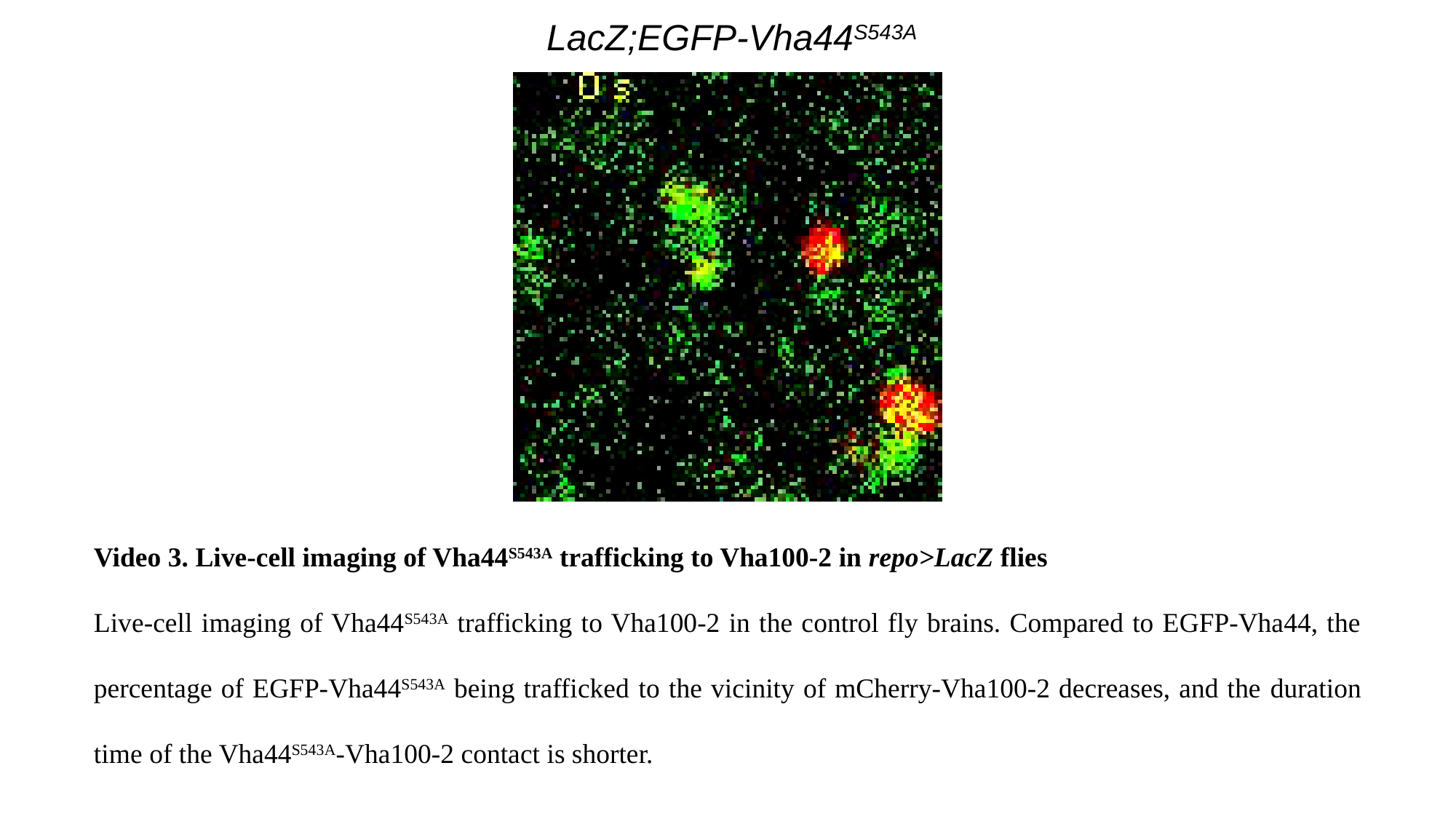

LacZ;EGFP-Vha44^S543A^
Video 3. Live-cell imaging of Vha44^S543A^ trafficking to Vha100-2 in repo>LacZ flies
Live-cell imaging of Vha44^S543A^ trafficking to Vha100-2 in the control fly brains. Compared to EGFP-Vha44, the percentage of EGFP-Vha44^S543A^ being trafficked to the vicinity of mCherry-Vha100-2 decreases, and the duration time of the Vha44^S543A^-Vha100-2 contact is shorter.

### Slide 4
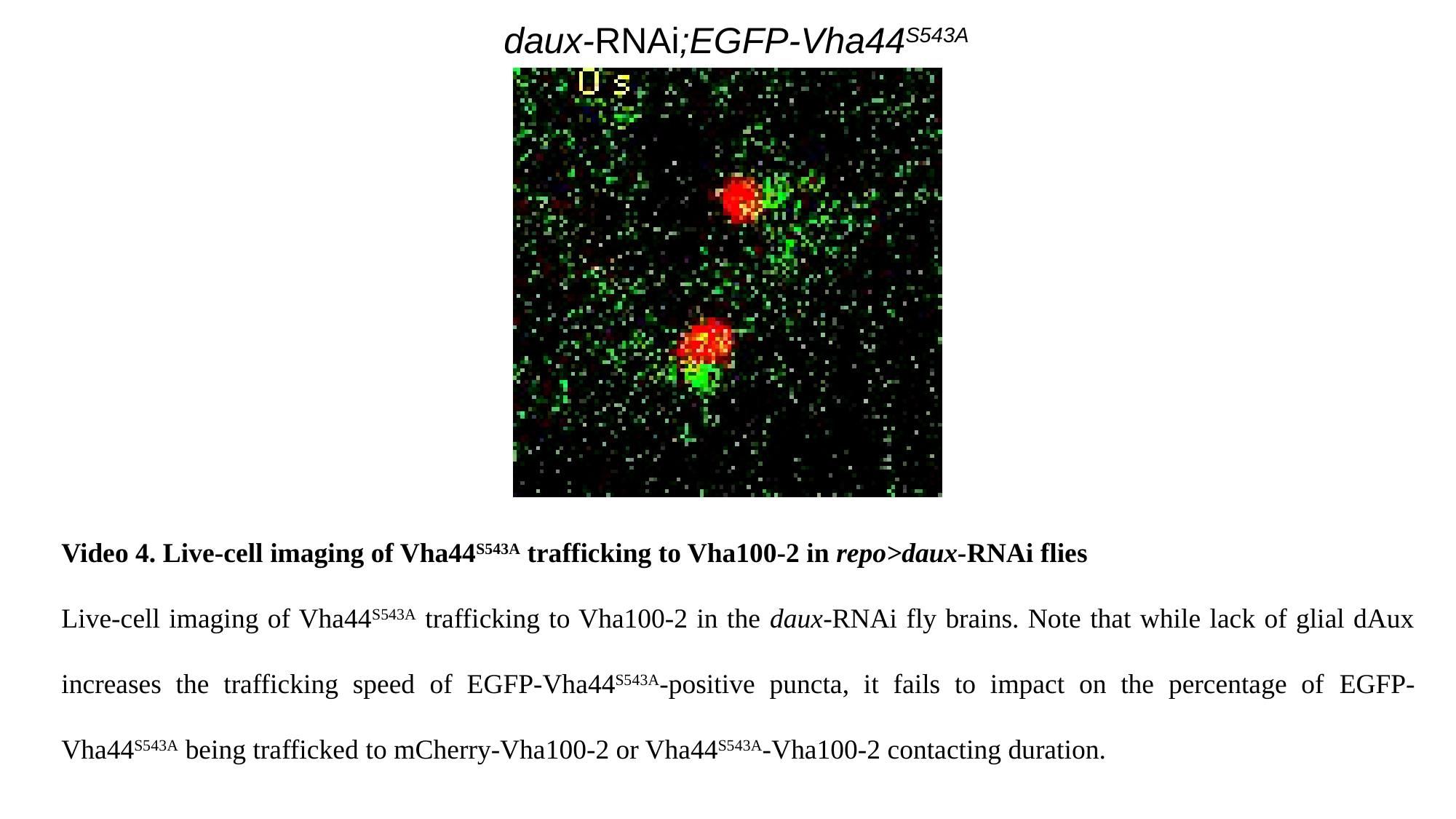

daux-RNAi;EGFP-Vha44^S543A^
Video 4. Live-cell imaging of Vha44^S543A^ trafficking to Vha100-2 in repo>daux-RNAi flies
Live-cell imaging of Vha44^S543A^ trafficking to Vha100-2 in the daux-RNAi fly brains. Note that while lack of glial dAux increases the trafficking speed of EGFP-Vha44^S543A^-positive puncta, it fails to impact on the percentage of EGFP-Vha44^S543A^ being trafficked to mCherry-Vha100-2 or Vha44^S543A^-Vha100-2 contacting duration.
